# Chromatin-associated RNA Dictates the ecDNA Interactome in the Nucleus

**DOI:** 10.1101/2023.07.27.550855

**Authors:** Zhengyu Liang, Collin Gilbreath, Wenyue Liu, Yan Wang, Michael Q. Zhang, Dong-Er Zhang, Sihan Wu, Xiang-Dong Fu

## Abstract

Extrachromosomal DNA (ecDNA) promotes cancer by driving copy number heterogeneity and amplifying oncogenes along with functional enhancers. More recent studies suggest two additional mechanisms for further enhancing their oncogenic potential, one via forming ecDNA hubs to augment oncogene expression ^1^ and the other through acting as portable enhancers to *trans-* activate target genes ^2^. However, it has remained entirely elusive about how ecDNA explores the three-dimensional space of the nucleus and whether different ecDNA have distinct interacting mechanisms. Here, by profiling the DNA-DNA and DNA-RNA interactomes in tumor cells harboring different types of ecDNAs in comparison with similarly amplified homogenously staining regions (HSRs) in the chromosome, we show that specific ecDNA interactome is dictated by ecDNA-borne nascent RNA. We demonstrate that the ecDNA co-amplifying *PVT1* and *MYC* utilize nascent noncoding *PVT1* transcripts to mediate specific *trans*-activation of both ecDNA and chromosomal genes. In contrast, the ecDNA amplifying *EGFR* is weak in this property because of more efficient splicing to remove chromatin-associated nascent RNA. These findings reveal a noncoding RNA-orchestrated program hijacked by cancer cells to enhance the functional impact of amplified oncogenes and associated regulatory elements.

## INTRODUCTION

Each mammalian chromosome occupies a distinct nuclear domain excluded from adjacent chromosome-occupied areas, known as the chromosome territory ^3,4^. Inside a territory, sub-megabase-scaled chromatins frequently form statistically interacting units, called topologically associating domains (TADs), to establish *cis*-regulatory networks ^5–7^. In addition, neighboring chromosomes may contact each other *in trans* ^8^, and non-homologous chromosome interactions can be achieved by translocation frequently detected in cancer ^9^.

Extrachromosomal DNA (ecDNA) has recently emerged as a prevalent driver of cancer ^10,11^. Besides amplifying specific oncogenes ^12–14^, ecDNA appear to have acquired additional mechanisms to drive tumorigenesis after escaping the chromosomal constraint by either forming transcription-promoting clusters termed ecDNA hubs to further enhance oncogene expression ^1,15^ or serving as mobile enhancers to stimulate target gene expression ^2,16^. However, these concepts are still under debate, as genomic targeting of ecDNA has not been disambiguated by comparison with the chromosomally amplified counterpart, such as the homogeneously staining region (HSR). And a separate super-resolution imaging study even fails to detect ecDNA hubs in glioblastoma ^17^.

The present study aims to address these outstanding problems through a systematic comparison between ecDNA and HSR interactomes at both DNA and RNA levels through functional perturbation for mechanistic dissection, revealing that specific ecDNA may amplify their oncogenic potential via the action of ecDNA-associated noncoding RNA.

## RESULTS

### Comparison between ecDNA and HSR in colon cancer cells

A previous Hi-C study suggests the ability of ecDNA to broadly target genomic loci ^2^; however, the data do not differentiate the contribution of increased gene copy number versus elevated ecDNA mobility, which would benefit from a comparison with similar focal amplification as HSR. To address this important question, we took advantage of colorectal cancer cell lines COLO320DM and COLO320HSR derived from the same patient, both of which carry amplified DNA segments containing *MYC* oncogene as determined by whole-genome sequencing (WGS, Extended Data Figure 1a-c). We validated that COLO320DM and COLO320HSR cells respectively harboring ecDNA and HSR by *MYC*-labeled DNA fluorescence *in situ* hybridization (FISH), as indicated by scattered foci versus focal FISH signals on metaphase nuclei (Figure 1a).

**Figure 1.**
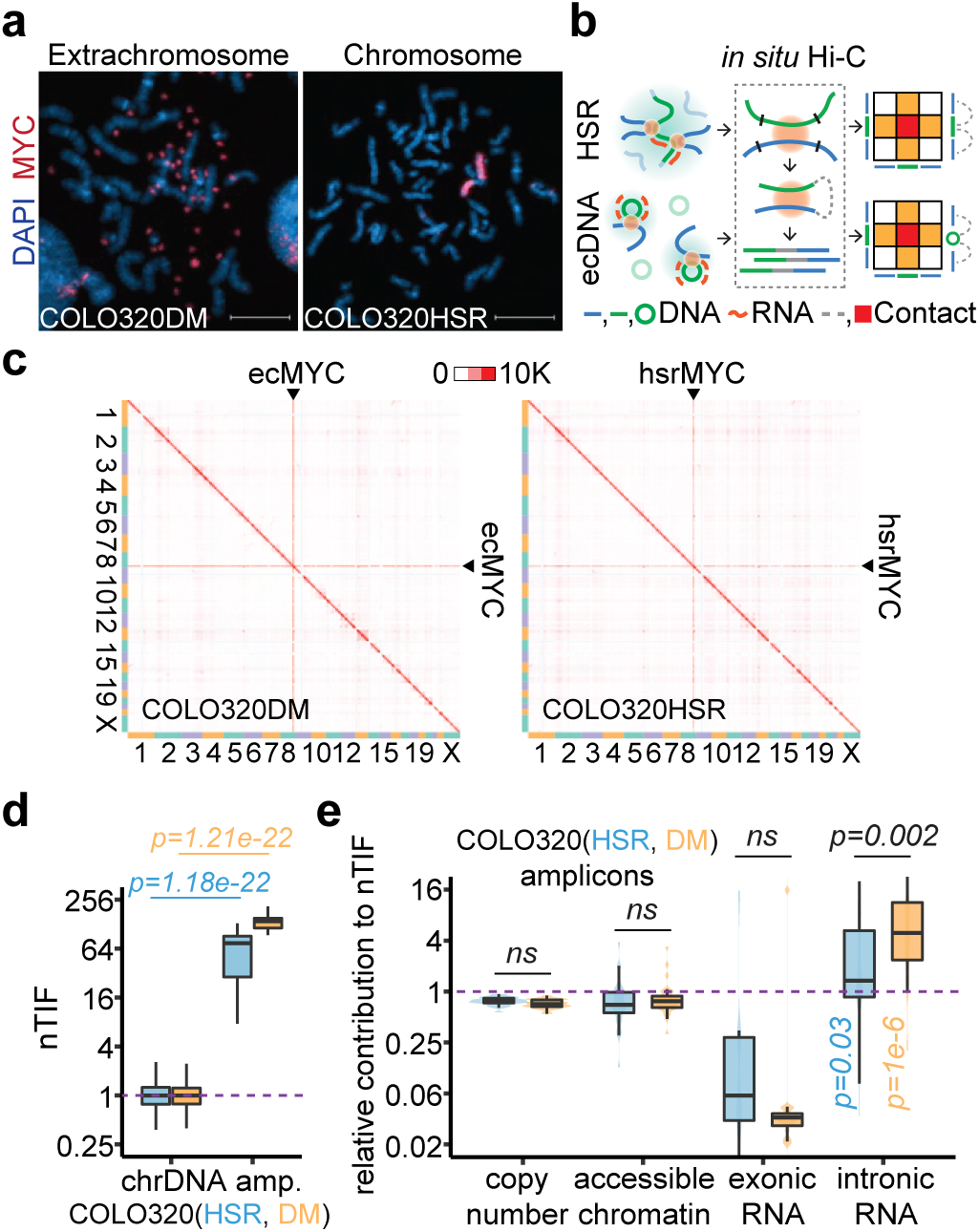
Elevated chromosomal interactions of ecDNA were enriched for unspliced RNA. **(a)** Representative *MYC* DNA FISH images showing ecDNA and HSR in metaphase chromosome spreads. Scale bars, 10 μm. **(b)** Schematic of *in situ* Hi-C assay detecting chromatin interactions of HSR or ecDNA. Amplicon chromatins from COLO320HSR cells (HSR, green line) or COLO320DM cells (ecDNA, green circle) were fragmented by restriction enzyme (black line) and ligated to chromosomal DNA (blue line) in proximity, resulting as interacting-frequency based contact heatmap (colored square). **(c)** *In situ* Hi-C contact heatmap for COLO320DM or COLO320HSR in the order of mapping position (chromosome 1 to 22, X) with ecMYC or hsrMYC labeled. Scale bars, 10,000 interacting reads. **(d)** Normalized *trans-*interaction frequency (nTIF) for DNA bins (50 kb resolution). nTIF of chromosomal DNA was scaled with the median of 1 (dotted line), and relative nTIF of hsrMYC (blue) and ecMYC (orange) were represented in the boxplot (One-sided Wilcoxon Rank Sum Test). **(e)** Normalization power of copy number (measured by WGS), accessible chromatin (measured by ATAC-seq), exonic RNA (measured by RNA-seq-derived exonic reads), or intronic RNA (measured by RNA-seq-derived intronic reads) for each binned nTIF. Normalization power of chromosomal DNA was scaled to the median of 1 (dotted line), and relative contribution of hsrMYC (blue) and ecMYC (orange) was shown in the boxplot (One-sided Wilcoxon Rank Sum Test. ns: not significant).

We next performed *in situ* Hi-C ^18,19^ on this pair of cell lines to map their 3D genomic interactions (Figure 1b). Using highly reproducible Hi-C data (Extended Data Figure 1d), we found that both ecDNA- and HSR-amplified regions exhibited dramatically increased *trans-*interactions (Figure 1c, Extended Figure 1e). Thus, the elevated *trans*-interactions could not be entirely attributed to the mobility of ecDNA, contrary to the original conclusion ^2^. To aid in quantitative analysis, we took a previous approach to calculate the normalized *trans*-chromosomal interaction frequency (nTIF) ^2^, and as expected, both ecDNA- and HSR-amplified regions showed much higher nTIF values compared to typical chromosomal regions (Figure 1d, Extended Data Figure 1f). This observation raises the question of which chromatin feature(s) contributes to such increased *trans*-interactions and, more interestingly, which feature(s) may differentiate between ecDNA and HSR.

We approached these questions by dividing the human genome into 1-kb bins and calculating each bin-associated amplified factor, such as DNA copy number (based on WGS), chromatin accessibility (determined by ATAC-seq), and the state of transcription (measured by RNA-seq) on ecDNA or HSR, for both amplicons and all chromosomal regions (Extended Data Figure 1g). We next normalized the corresponding nTIF values with these amplified factors to obtain the adjusted nTIF values (nTIF^adj^), whose inverted values would then reflect the power of individual amplified factors in contributing to enhanced *trans*-interactions. Finally, we determined the relative power of amplified regions relative to normal chromosomal regions and asked which chromatin feature(s) might differentiate between ecDNA and HSR. This analysis revealed that intronic RNA signals had the highest power to differentiate between ecDNA and HSR (Figure 1e). This finding points to the critical contribution of chromatin-associated nascent RNA to a potential functional distinction between ecDNA and HSR.

### ecDNA mobility manifested by ecDNA-borne nascent RNA

To determine nascent RNA-mediated ecDNA interactome, we performed *in situ* global RNA interactions with DNA by deep sequencing (GRID-seq), which we developed earlier to profile chromatin-associated RNAs (caRNAs) ^20,21^. Following the established protocol ^20^, we generated highly reproducible GRID-seq data on COLO320HSR and COLO320DM cells (Extended Data Figure 2), showing that most caRNAs were nascent RNAs that contain introns as compared to the RNA-seq data. Still, both were comparable between ecDNA and HSR (Figure 2b). Because only 25% of DNA-RNA pairs were uniquely mapped, we took a multiple alignment strategy (Methods) to increase the detection sensitivity (Extended Data Figure 3a-d). We normalized RNA expression to reads per kb per million (RPKM). As previously described ^21^, we built a background model by assuming that most protein-coding RNAs (pcRNAs) randomly interact with DNAs after being released from their sites of transcription (Extended Data Figure 3e), thus helping define specific *trans*-interactions.

**Figure 2.**
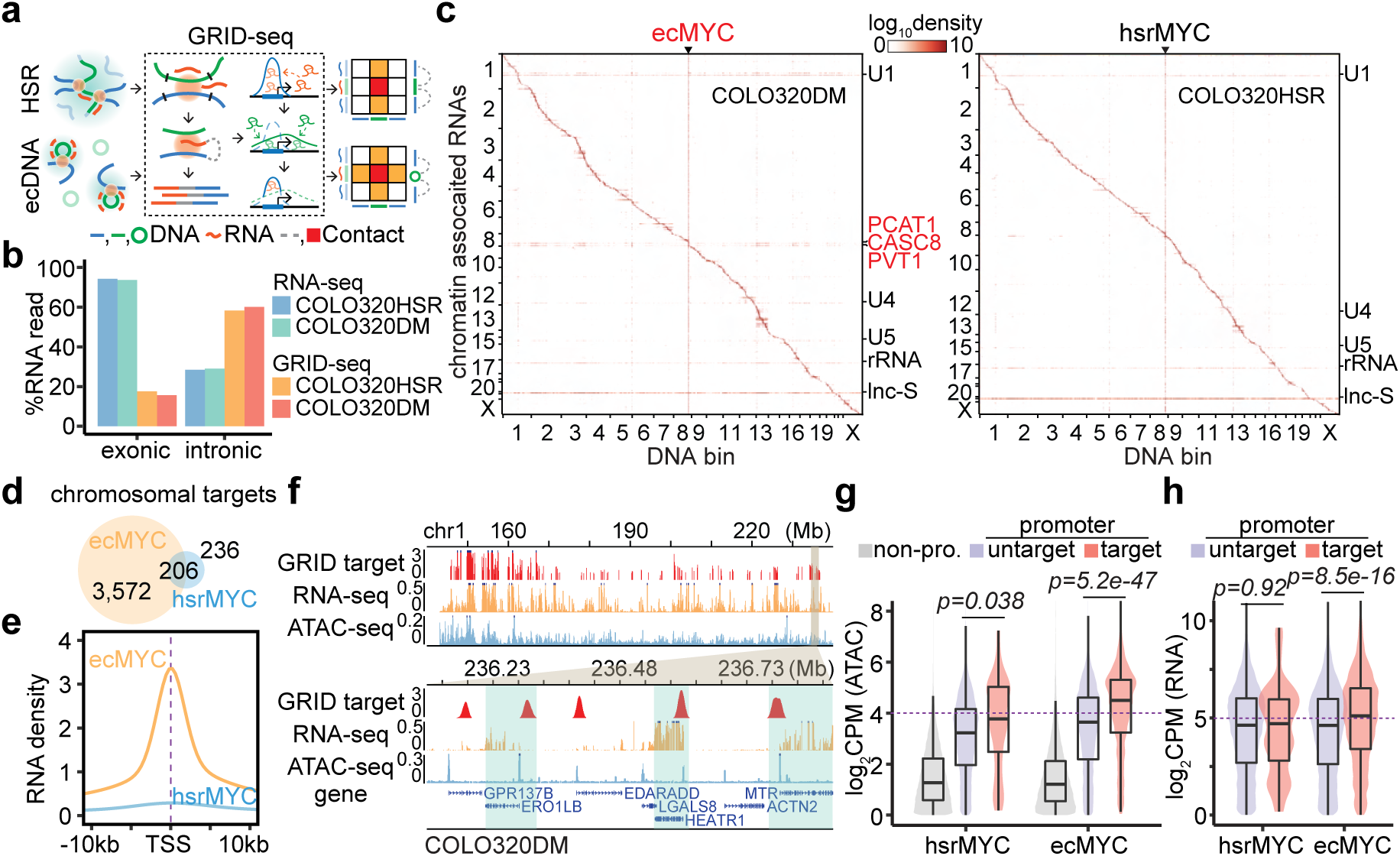
ecDNA acted as a mobile enhancer targeting highly expressed chromosomal genes in GRID-seq detection. **(a)** Schematic of GRID-seq detecting chromatin-enriched RNA with DNA interactions of HSR or ecDNA. HSR (green line) or ecDNA (green circle) derived RNAs (red line) were ligated to fragmented chromosomal DNA (blue line) in proximity and converted into DNA libraries for strand-specific paired-tag detection. *Trans-*chromosomal interacted protein-coding RNAs (green) were used to build the background model of random interactions. Chromatin-interacting RNAs (red) above the background (green dotted line) were calculated as relative enrichment to construct the RNA-DNA contact heatmap (colored square). **(b)** Quantification of RNA signal annotated by exonic or intronic reads in the detection of RNA-seq or GRID-seq. **(c)** Heatmap showing chromatin-enriched RNAs interacted across the whole genome in the order of mapping position, with rows for chromatin-enriched RNAs and columns for DNA bins in 2-Mb resolution. ecMYC or hsrMYC were labeled on top, and major *trans-*chromosomal interacting RNAs were labeled on the right (ecDNA-derived RNAs in red). Scale bars, relative enrichment in log-10 scale. **(d)** A Venn diagram showing the number of chromosomal targets interacted by ecMYC or hsrMYC-derived RNAs in COLO320DM or COLO320HSR cells. **(e)** Meta-analysis of ecMYC or hsrMYC targeted chromosome regions relative to annotated TSS. Interacting RNAs were weighted for ecMYC (orange) and hsrMYC (blue). **(f)** Representative genomic loci showing chromosomal interacting probability of ecDNA-borne RNAs (log-10 relative fold enrichment in GRID-seq, red), chromosomal genes expression (count-per-million, CPM, in RNA-seq, orange), and chromatin accessibility (CPM in ATAC-seq, blue) for COLO320DM cells. Top, overview. Bottom, zoom-in view. **(g)** Quantification of chromatin accessibility by ATAC-seq. Grey, non-promoter regions; Red, hsrMYC or ecMYC targeted promoters; Purple, non-targeted promoters. P-values were calculated by one-sided Wilcoxon Rank Sum Test. **(h)** Quantification of chromosomal genes expression by RNA-seq. Red, genes with promoters targeted by hsrMYC or ecMYC; Purple, genes with promoters untargeted. P-values were calculated by one-sided Wilcoxon Rank Sum Test.

After these data treatments, we first noted that the amplified genes produced high levels of RNAs detected by both RNA-seq and GRID-seq in both COLO320DM and COLO320HSR cells (Extended Figure 4a,b). In agreement with the previous findings ^21^, most caRNAs frequently interacted with local chromatins at or near the originated gene loci (i.e. within the distance of 10 kb), as exemplified by the pcRNA from the *TBC1D4* gene (Extended Data Figure 4c, top). Thus, most RNA-DNA interactions were visible along the diagonal line on the contact heatmap in both COLO320DM and COLO320HSR cells (Figure 2c). The *MYC-*amplified DNA region exhibited a massive increase in interactions with RNAs genome-wide, as indicated by the vertical line on the contact heatmap, regardless of ecDNA or HSR, indicating that this interaction pattern is primarily driven by DNA amplification as detected by Hi-C (see Figure 1c). In contrast, after background correction, ecDNA-borne RNAs in COLO320DM, but not HSR-derived RNAs in COLO320HSR, exhibited broad *trans*-interactions across the genome, showing a horizontal line on the contact heatmap, similar to the U1 spliceosomal RNA known to predominately engage in *trans*-interactions (Figure 2c, Extended Data Figure 4c, middle and bottom). Thus, the mobility of ecDNA is characterized by broad *trans*-interactions of ecDNA-borne nascent RNAs in COLO320DM cells, but not in COLO320HSR cells.

### ecDNA exploring open chromatin to activate target genes

To identify specific genes interrogated by trans-interactions, we utilized all HSR- or ecDNA-borne RNAs to identify their DNA targets by requiring 2-fold enrichment on individual 1-kb DNA bins over the background in COLO320HSR and COLO320DM cells (Extended Data Figure 4d). We next merged continuous DNA bins into single units, leading to the identification of 3,572 and 236 chromosomal targets for ecDNA- versus HSR-transcribed RNAs, with 206 shared targets (Figure 2d). The sharp difference in the number of targets again agrees with the much greater mobility of ecDNA compared to HSR. Although RNAs transcribed from HSR and ecDNA showed similar preference for gene promoters over randomly selected genomic segments (Extended Data Figure 4e), ecDNA-interacting sites exhibited a higher degree of enrichment at transcription start sites (TSSs, Figure 2e), demonstrating that ecDNA preferentially targets gene promoters both in quantity and quality.

We next asked whether ecDNA-engaged *trans-*interactions are linked to chromosomal activities and gene expression. We found that ecDNA-targeted sites were generally associated with accessible chromatin and active transcription in COLO320DM cells (Figure 2f). ecDNA-targeted promoters clearly exhibited higher accessibility than non-promoter or untargeted promoter regions (Figure 2g). In concordant, ecDNA-targeted genes showed higher transcriptional activities than untargeted genes, which is not the case for HSR-targeted genes (Figure 2h). Moreover, the H3K27ac signals on ecDNA, which reflect active enhancers, were significantly proportional to the propensity of those genomic regions to engage in *trans* DNA-DNA interactions detected by Hi-C (Extended Data Figure 4f,g). Together, these data show that ecDNA acts as a more active enhancer than HSR, despite similar H3K27ac profiles after correcting for DNA copy number variations (Extended Data Figure 4h).

### Different ecDNAs showing distinct targeting potential

The ecDNA interaction with chromatin may reflect a property of ecDNA in general; alternatively, it could be context-dependent with regard to the ecDNA sequence, copy number, or chromatin environment. To test these possibilities, we extended our analyses to another pair of similarly focal amplificed cancer cell lines derived from the same patient with glioblastoma, GBM39EC and GBM39HSR, which amplify *EGFR* as ecDNA with an averaged copy number of 86 or HSR with an average copy number of 91, which we validated by WGS and FISH (Figure 3a; Extended Data Figure 5a-c). In addition, GBM39EC cells also carry ecDNAs that modestly amplify *MYC*. As with the pair of colon cancer cells, we observed by *in situ* Hi-C that both GBM39EC and GBM39HSR exhibited extensive, yet similar, *trans* DNA-DNA interactions (Figure 3b and Extensive Data Figure 5d,e), as quantified by nTIF (Figure 3c), which roughly correspond to the amplified copy number for *EGFR* or *MYC*. Somewhat unexpectedly, we detected little difference in trans-interactions between GBM39EC and GBM39HSR, even after the adjustment based on copy number, chromatin accessibility, spliced RNA, and even nascent RNA (Figure 3d). We noted, however, that the *MYC* ecDNA appeared to show significant differences in intronic RNA-normalized *trans*-interactions in GBM39EC, thus resembling its behavior in COLO320DM cells.

**Figure 3.**
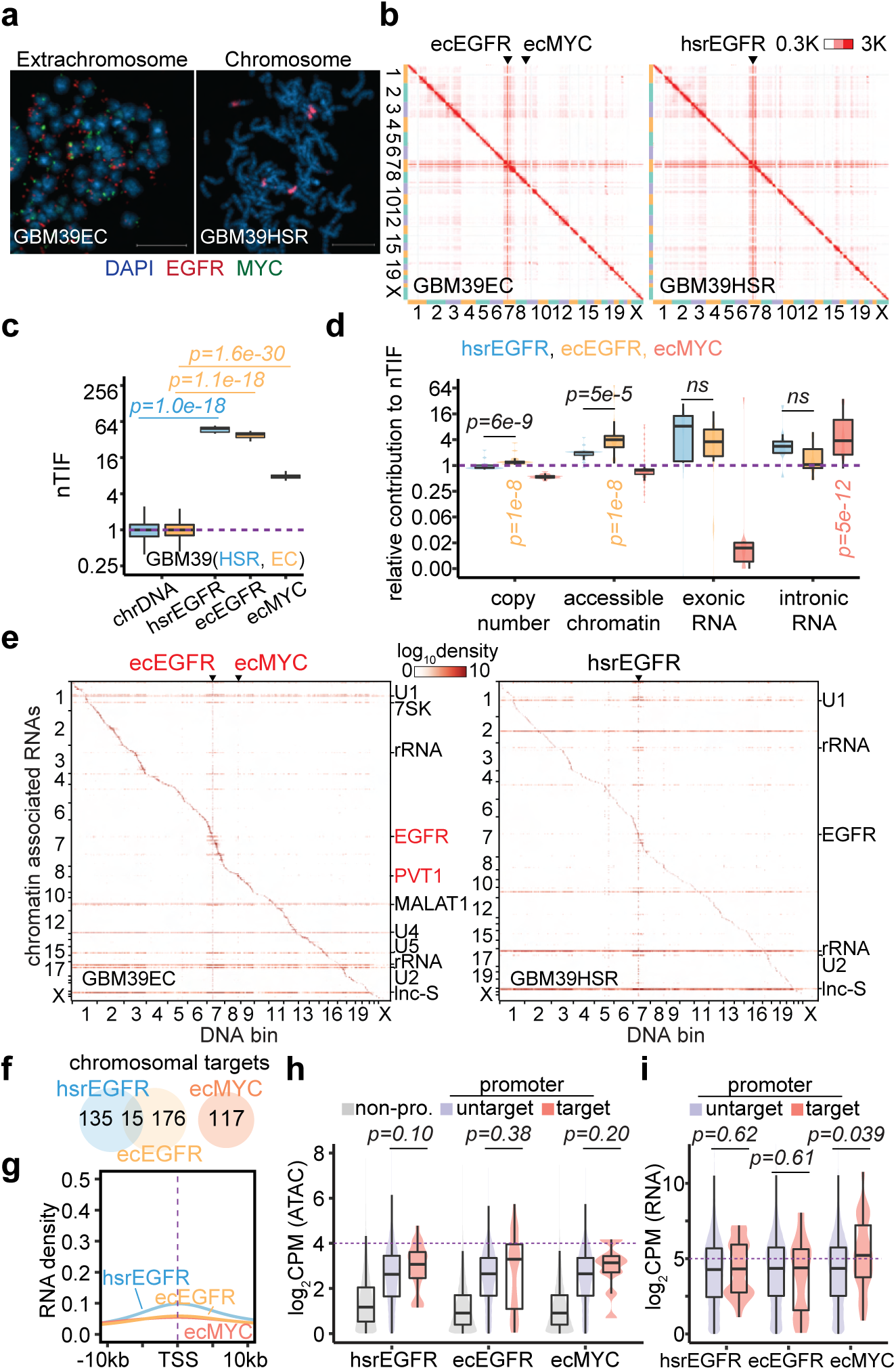
Limited mobile targeting of ecDNA in GBM39 cells. **(a)** Representative FISH images showing *EGFR* and *MYC* ecDNAs in GBM39EC and *EGFR* HSR in GBM39HSR cells at metaphase. Scale bars, 10 μm. **(b)** *In situ* Hi-C contact heatmap for GBM39EC and GBM39HSR cells in the order of mapping position (chromosome 1 to 22, X) with ecEGFR, ecMYC, and hsrEGFR labeled. Scale bars, 300 to 3,000 interacting reads. **(c)** Normalized *trans-*interaction frequency (nTIF) for DNA bins (50 kb resolution) presented similarly as in Figure 1d (one-sided Wilcoxon Rank Sum Test). **(d)** Normazliation power of copy number, accessible chromatin, exonic RNA, or intronic RNA for each binned nTIF as in Figure 1e (one-sided Wilcoxon Rank Sum Test; ns, not significant). **(e)** Heatmap showing chromatin-enriched RNAs interacted across the whole genome in the order of mapping position, with rows for chromatin-enriched RNAs and columns for DNA bins in 2-Mb resolution. ecEGFR, ecMYC, and hsrEGFR were labeled on top, and major *trans-*chromosomal interacting RNAs were labeled on the right (ecDNA-derived RNAs in red). Scale bars, 1og-10 relative enrichment. **(f)** A Venn diagram shows the number of chromosomal targets interacted by ecEGFR, ecMYC, or hsrEGFR-derived RNAs in GBM39EC or GBM39HSR cells. **(g)** Meta-analysis of ecEGFR, ecMYC, or hsrEGFR targeted chromosome regions relative to annotated TSS. Interacting RNAs were weighted for ecEGFR (orange), ecMYC (red), and hsrEGFR (blue). **(h)** Quantification of chromatin accessibility by ATAC-seq. Grey, non-promoter regions; Red, hsrEGFR, ecEGFR, or ecMYC targeted promoters; Purple, non-targeted promoters. P-values were calculated by one-sided Wilcoxon Rank Sum Test. **(i)** Quantification of chromosomal genes expression by RNA-seq. Red, genes with promoters targeted by hsrEGFR, ecEGFR, or ecMYC; Purple, genes with promoters untargeted. P-values were calculated by one-sided Wilcoxon Rank Sum Test.

We also performed GRID-seq on this pair of glioblastoma cell lines (Extended Data Figure 6). We detected a similar increase in the ability of amplified DNAs to interact with RNAs across the genome, regardless of ecDNA or HSR, as indicated by similar vertical lines in the DNA-RNA interaction heatmaps (Figure 3e). In contrast to the colon cancer cells that harbored high copy *MYC* ecDNA, we detected limited interactions of *EGFR*-derived caRNAs with genomic DNA, regardless of whether they were from ecDNA or HSR, as reflected by weak, if any, horizontal lines on the RNA-DNA interaction heatmaps as opposed to strong *trans*-acting spliceosomal RNAs (Figure 3e and Extended Data Figure 7a-c). We only identified 191 or 150 chromosomal targets for ecDNA or HSR-derived *EGFR* RNA, with only 15 targets in common. In comparison, despite modest amplification, we still scored 117 chromosomal targets for *MYC* ecDNA in GBM39EC cells (Figure 3f and Extended Data Figure 7d). These data suggest that *EGFR* ecDNA is engaged in *trans-*interactions with much low frequency in GBM39EC (Figure 3g and Extended Data Figure 7e). Additionally, the limited number of gene promoters targeted by *EGFR* ecDNA or HSR showed little difference in chromatin accessibility or transcriptional activities compared to untargeted promoters, whereas *MYC* ecDNA was still linked to more active gene promoters (Figure 3h,i). These data suggest that *EGFR* ecDNA is much more immobile than *MYC* ecDNA, suggesting that different ecDNAs have distinct *trans*-acting abilities, thus giving rise to ecDNA-specific interactomes in the nucleus.

### Noncoding *PVT1* RNA-mediated *MYC* ecDNA interactome

The effective association of RNAs with *trans*-interactions motivated us to further examine *trans*-acting caRNAs detected by GRID-seq. We separately determined the density of intronic reads, which reflects nascent unspliced RNAs, and the density of exonic reads, which includes both nascent unspliced and spliced nuclear-retained RNAs (Figure 4a). By caleculating the density ratio of intronic reads versus exonic reads for individual RNAs expressed from the amplicon, we found that in COLO320 cells, most ecDNA-derived *trans*-acting RNAs were unspliced RNAs, which were much reduced with HSR-derived *trans*-acting RNAs (Figure 4b). After normalization against background *trans*-acting RNAs (Methods), virtually the majority of *trans*-acting RNAs above the background were unspliced RNAs in ecDNA but not in HSR (Figure 4c and Extended Data Figure 8a). In GBM39 cells, *trans*-acting RNAs from the *MYC* amplicon were similar to those in COLO320 cells, despite modest gene amplification, and in contrast, most *trans*-acting RNAs from the *EGFR* amplicon were spliced RNAs, regardless of their production from ecDNA or HSR (Figure 4d and Extended Data Figure 8b) and most signals vanished after background correction (Extended Data Figure 8c). These results suggest that the behavior of *trans*-acting RNAs was amplicon-specific, not cell type-specific.

**Figure 4.**
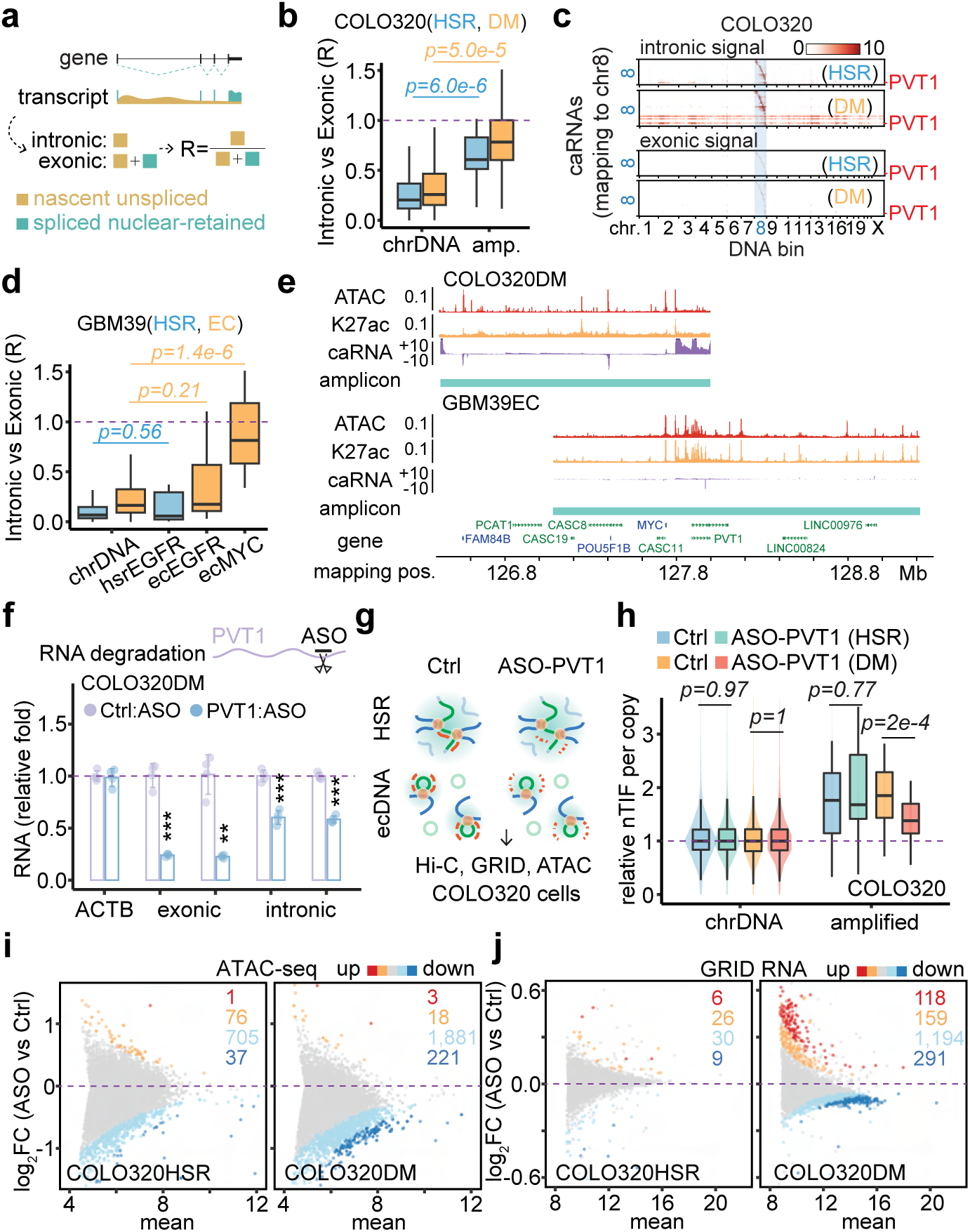
ecDNA-borne nascent noncoding RNAs mediate ecDNA interactome and transcriptional regulation *in trans*. **(a)** Schematic of nascent unspliced RNA signal and spliced nuclear-retained RNA signal. **(b)** Quantification of caRNA signal as intronic vs exonic ratio for COLO320HSR (blue) and COLO320DM (orange) cells. P-values were calculated by one-sided Wilcoxon Rank Sum Test. **(c)** Heatmaps of hsrMYC or ecMYC-derived RNAs interacting with the genome in the order of mapping position. Top heatmaps, intronic caRNA derived. Bottom heatmaps, exonic caRNA-derived. Scale bars, log-10 relative enrichment. **(d)** Quantification of caRNA signal as intronic vs exonic ratio for cells GBM39HSR (blue) and GBM39EC (orange). P-values were calculated by one-sided Wilcoxon Rank Sum Test. **(e)** Genomic tracks showing ATAC-seq, H3K27ac ChIP-seq, and GRID-seq-derived RNA (strand-specific, in two scaled views) signals normalized to CPM in COLO320DM cells. Amplicons were determined by AmpliconArchitect software. **(f)** qRT-PCR for relative RNA levels of internal control *ACTB*, exonic and intronic *PVT1*, comparing *PVT1* knockdown to control in COLO320DM cells (two-sided Student’s t-test). **(g)** Schematic of *in situ* Hi-C, GRID-seq, and ATAC-seq assays in COLO320HSR (top) and COLO320DM (bottom) cells treated by control or *PVT1* ASO knockdown. **(h)** Quantification of copy number normalized nTIF for chromosomal DNA bins (scaled to median of 1) and relative nTIF of HSR or ecDNA treated by control or *PVT1* knockdown (one-sided Wilcoxon Rank Sum Test). **(i)** Scatter plot of log-2 fold change of normalized ATAC-seq signal upon *PVT1* knockdown against the control sample. Highlighted are promoters with signal significantly increased (red, FDR<0.01; orange, FDR<0.05) or decreased (dark blue, FDR<0.01; light blue, FDR<0.05). **(j)** Scatter plot of log-2 fold change of normalized GRID-seq-derived RNA expression upon *PVT1* knockdown against the control sample. Highlighted are genes with expression significantly increased (red, FDR<0.01; orange, FDR<0.05) or decreased (dark blue, FDR<0.01; light blue, FDR<0.05).

The noncoding *PVT1* transcripts are the major RNA transcribed from *MYC* ecDNA in both COLO320DM and GBM39EC cells (Figure 4e, Extended Data Figure 8d). This prompted us to investigate whether this ecDNA-borne noncoding RNA plays a direct role in mediating ecDNA targeting in the genome. To this end, we used a previously validated antisense oligo (ASO), which targets exon 2 of the *PVT1* gene ^22^, to knock down the nascent *PVT1* transcript. This ASO successfully decreased the levels of exonic and intronic reads of *PVT1* (Figure 4f). Next, we applied this ASO-*PVT1* in COLO320HSR and COLO320DM cells and performed *in situ* Hi-C, GRID-seq, and ATAC-seq to determine the potential *PVT1* RNA-mediated function of ecDNA (Figure 4g). We found that, upon *PVT1* knockdown, *MYC-PVT1* co-amplified regions still exhibited elevated *trans*-interactions detected by Hi-C and GRID-seq across the whole genome in COLO320HSR and COLO320DM cells (Extended Data Figure 8e,f). However, quantification based on DNA copy number-adjusted nTIF showed that *PVT1* knockdown significantly decreased the *trans*-interaction frequency of *MYC* ecDNA in COLO320DM cells, but not *MYC* HSR in COLO320HSR cells (Figure 4h). Of note, ASO-*PVT1* did not change the copy number of *MYC* ecDNA in COLO320DM (Extended Data Figure 8g). These data provide evidence for *PVT1*-mediated ecDNA mobility.

Accompanying the attenuation of *MYC* ecDNA *trans*-interactions were significant changes in global chromatin accessibility and transcriptome. By ATAC-seq, we detected reduced promoter accessibility upon *PVT1* knockdown, which was more pronounced in COLO320DM cells compared to COLO320HSR cells (Figure 4i). Taking advantage of the ability of GRID-seq to detect nascent RNA production similar to GRO-seq ^21^, we found that *PVT1* knockdown also attenuated the transcriptome in COLO320DM cells, with 1,485 down-regulated genes compared to 277 up-regulated ones. In contrast, *PVT1* knockdown only slightly affected the transcriptome in COLO320HSR cells, with 39 and 32 down- or up-regulated genes detected (Figure 4j). Of note, the effect size was modest, likely because the set of ecDNA molecules could only interact with a limited number of targets in a given cell, and the target set likely varies from cell to cell, thus diminishing the functional impact measured on a population of cells. In any case, genes from *MYC* ecDNA-targeted promoters appear more down-regulated compared to those from non-targeted promoters upon *PVT1* knockdown in COLO320DM cells, but not in COLO320HSR cells (Extended Data Figure 8h). Finally, we observed a significant concordance between *PVT1* knockdown-induced reduction in promoter accessibility and transcription in COLO320DM, but not in COLO320HSR cells (Extended Data Figure 8i). Collectively, these data suggest that ecDNA-borne nascent *PVT1* mediates genomic targeting of *MYC* ecDNA to activate target gene expression.

### Coordination between cDNA hub formation and genomic targeting

*MYC* ecDNA molecules were found to form clusters termed hubs in COLO320DM cells to enhance the expression of the ecDNA-borne oncogene(s) ^1^. It remains unclear whether such intermolecular interactions of ecDNA coordinate or compete with the ability of ecDNA to interact in *trans* to activate chromosomal gene transcription. Using FISH on interphase nuclei, we recognized at least three types of ecDNA hubs: 1) The majority of ecDNA particles formed one to a few micron-size superstructures, such as *MYC* ecDNAs in COLO320DM cells. We named these ecDNA speckles; 2) Most ecDNAs were scattered in the nucleus, with only a few ecDNA clusters in proximity, such as *EGFR* and *MYC* ecDNAs in GBM39EC cells. We called it ecDNA minim; 3) Two or more species of ecDNAs intertwined to form mosaic ecDNA speckles besides ecDNA minim, as exemplified with *MYC* and *FGFR2* ecDNAs in SNU16 cells (Figure 5a). In general, different types of ecDNAs appear to have distinct potential to aggregate.

**Figure 5.**
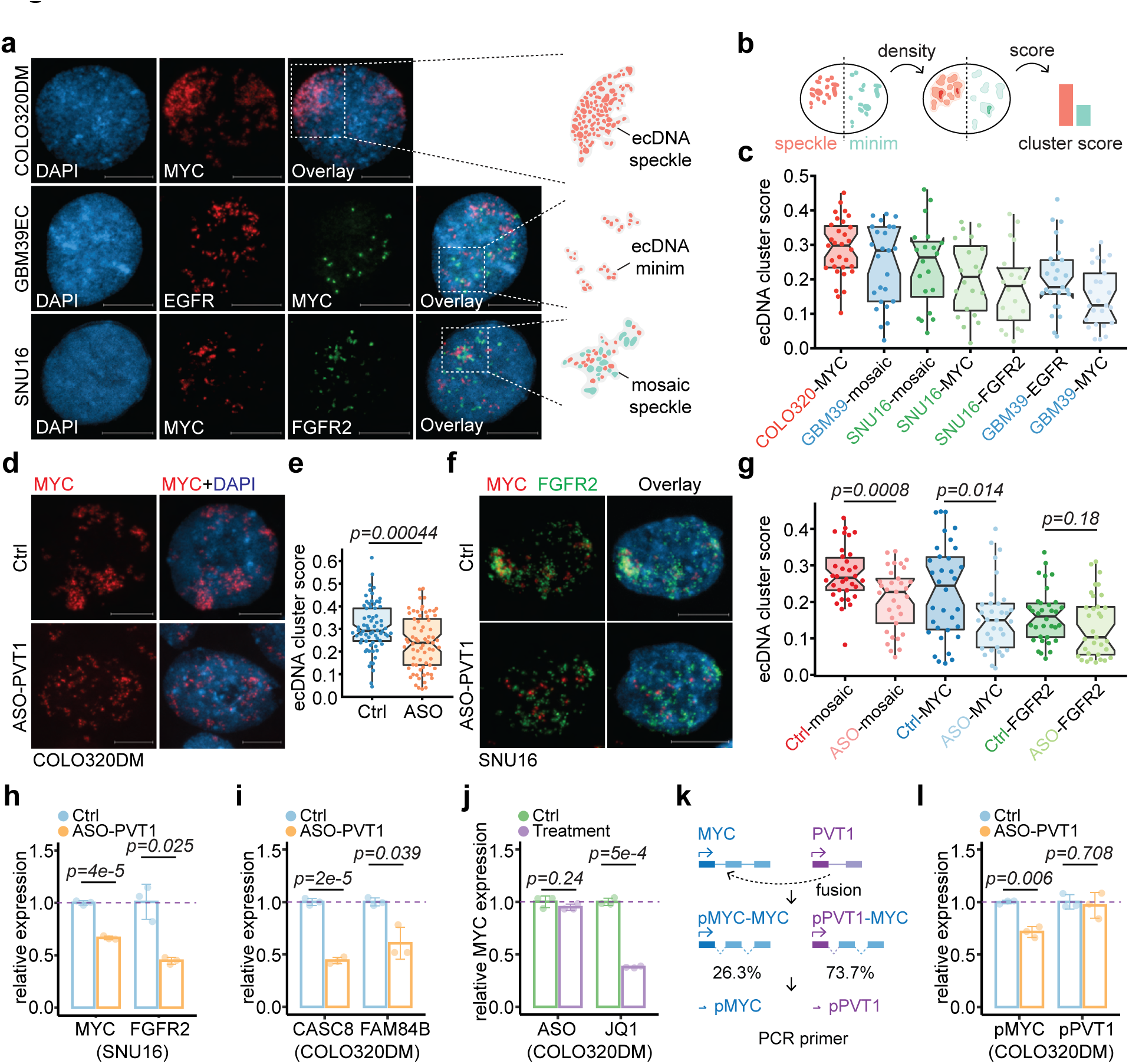
ecDNA-borne RNAs mediated ecDNA hub formation. (a) Representative FISH images in interphase nuclei showing ecMYC in COLO320DM, ecEGFR and ecMYC in GBM39EC, and ecMYC and ecFGFR2 in SNU16 cells. Scale bars, 10 μm. (b) Schematic of probability density-based method as ecDNA clustering score for both ecDNA speckle (red) and ecDNA minim (green). (c) Scored interphase ecDNA clustering in cancer cell lines. **(d-e)** Representative FISH images of interphase ecMYC treated by control or *PVT1* knockdown (d) and scored (e) in COLO320DM cells. **(f-g)** Representative FISH images of interphase ecMYC (green) and ecFGFR2 (red) treated by control or *PVT1* knockdown (f) and scored (g) in SNU16 cells. **(h)** qRT-PCR for relative *MYC* and *FGFR2* RNA levels in the treatment of *PVT1* knockdown against control in SNU16 cells (two-sided Student’s t-test). **(i)** qRT-PCR for relative *CASC8* and *FAM84B* RNA levels in the treatment of *PVT1* knockdown against control in COLO320DM cells (two-sided Student’s t-test). **(j)** qRT-PCR for relative *MYC* RNA levels in the treatment of *PVT1* knockdown against control (left) or JQ1 against control (right) in COLO320DM cells (two-sided Student’s t-test). **(k)** Schematic of *MYC* exon driven by *MYC* promoter (blue) and *PVT1* promoter (purple) and primers targeting wild-type (pMYC) and fusion *PVT1*-*MYC* (pPVT1). **(l)** qRT-PCR for relative *MYC* or *PVT1* promoter-derived RNA levels in the treatment of *PVT1* knockdown against control in COLO320DM cells (two-sided Student’s t-test).

To quantitatively measure the degree of ecDNA aggregation, we developed an ecDNA cluster score to capture the likelihood of ecDNA aggregation by using the probability density function-based Gaussian fitting and thresholding (Methods). For example, the score will be higher if highly amplified ecDNA particles form a few large ecDNA speckles; in contrast, the score will be lower if only a few ecDNAs form ecDNA minims in a background of many randomly distributed ecDNA molecules (Figure 5b). Therefore, our ecDNA cluster score is weighed by the probability of ecDNA aggregation, ecDNA hub size, and ecDNA hub number. Based on a series of computationally simulated images, we were able to show that the ecDNA cluster scoring method matches our domain knowledge and outperforms the previous approach based on autocorrelation (Extended Data Figure 9a,b).

Using the newly developed algorithm, we found that *MYC* ecDNA in COLO320DM cells had a significantly higher ecDNA cluster score than *EGFR* ecDNA and *MYC* ecDNA in GBM39EC cells, the latter of which might result from a lower copy number relative to that in COLO320DM (Figure 5c). Most strikingly, ASO-mediated knockdown of the ecDNA-borne *PVT1* transcripts significantly dispersed ecDNA speckles into ecDNA minims in both COLO320DM and SNU16 cells (Figure 5d-g). Notably, in SNU16 cells, mosaic ecDNA speckles were more profoundly affected by *PVT1* knockdown with significance, but not ecDNA minims such as *FGFR2* ecDNA, which does not contain *PVT1*. Although *MYC* ecDNA hubs were significantly dispersed, the statistical significance was primarily driven by the decrease of higher cluster scores, not lower ones (Figure 5g), suggesting that ecDNA speckle formation may be more dependent on the transcription of ecDNA-borne lncRNA. Collectively, these data show that ecDNA mobile targeting and hub formation are ordinated rather than in competition, and importantly, both are regulated by ecDNA-borne nascent lncRNA.

### Coordinated impact on ecDNA clustering and transcription

Finally, to generalize the functional impact of ecDNA-mediated *trans*-interactions, we performed ASO-*PVT1* knockdown in SNU16 cells where *MYC* and *FGFR2* ecDNAs were shown to hitchhike each other’s enhancer to promote oncogene expression ^1^. Aligned with this regulatory paradigm, we also observed that, upon *PVT1* knockdown and mosaic ecDNA speckle dispersion in SNU16 cells, *MYC* and *FGFR2* expression were both significantly decreased (Figure 5h). However, to our surprise, although the expression of two major *MYC* ecDNA-borne genes, *CASC8* and *FAM84B*, was significantly down-regulated after *MYC* ecDNA speckle dispersion by *PVT1* knockdown in COLO320DM cells (Figure 5i), the expression of *MYC* itself was unaffected by ASO-*PVT1* treatment. This observation contrasts intriguingly with a previous study ^1^, in which *MYC* ecDNA hub dispersion by BET bromodomain inhibitor JQ1 treatment led to the down-regulation of *MYC* expression (Figure 5j).

To address this discrepancy, we performed Hi-C, suggesting that, in COLO320DM cells, 73.7% of the *MYC* gene was driven by the *PVT1* promoter, resulting from the fusion of the *PVT1* promoter to *MYC* exon 2-3 (Figure 5k, Extended Data Figure 9c-e), which is consistent with a previous estimation based on fusion transcripts detected ^1^. Using region-specific primers for qPCR, we found that while the activity of the wild-type *MYC* promoter was decreased upon *PVT1* knockdown, in agreement with the enhancer hitchhiking model via ecDNA speckle, the activity of the *PVT1* promoter was unaffected by ASO-*PVT1* (Figure 5l). Therefore, the majority of *MYC* transcripts remained unchanged because they were driven by the ASO-insensitive *PVT1* promoter, even after *MYC* ecDNA speckle dispersion. These data suggest gene fusion-mediated promoter hijacking may secure oncogene output even when enhancer input is disrupted.

## DISCUSSION

ecDNA is an emerging hallmark of human cancer, whose high copy number, hyper-accessible chromatin, and random segregation enable tumors to evolve rapidly and develop resistance to therapy, thereby contributing to shorter survival for patients. In this study, we show that ecDNA interactome is mediated by ecDNA-borne nascent noncoding RNA, including *trans* ecDNA-chromosome interactions and ecDNA-ecDNA aggregation, which enables enhancer hitchhiking to promote gene transcription, revealing another layer of complexity of ecDNA biology in the cancer transcriptome.

Genomic targeting of mobile ecDNA was previously proposed through chromatin interaction analysis with paired-end tag (ChIA-PET) sequencing, coupled with RNA polymerase II pull-down enrichment for interacting chromatins ^2^. In addition, artificially synthesized circular DNAs containing enhancer sequences from ecDNA were used to support that ecDNA mimicry was capable of up-regulating chromosomal gene expression ^2^. However, it has been unclear whether such chromatin enrichment before interaction analysis would introduce a bias toward enhancer-promoter interaction, and to what extent artificial DNA circles recapitulate the biological context of ecDNA-chromosome interaction and function as previously discussed^16^. In our study, we opted to apply unbiased technologies, including Hi-C and GRID-seq, to two model systems with similarly focal amplified controls. Our data reveal that *MYC* ecDNA is a *de facto* mobile enhancer in COLO320DM cells, in which the ecDNA-borne nascent noncoding RNA *PVT1* bridges ecDNA’s enhancer to interact with the promoters of specific chromosomal genes. Such a mechanism may also explain why the *EGFR* ecDNA in GBM39EC is less mobile because it expresses a lower abundance of nascent noncoding RNA.

We note that the effect size of *PVT1* knockdown on chromosomal gene transcription is quite modest. This is likely due to the limited number of targeted genes in a single cell; thus, specific transcriptional impact is diluted from a population of cells. In COLO320DM cells, even though the *MYC* ecDNA was estimated to have 195 copies, we identified 3,572 chromosomal targets by unbiased GRID-seq. Therefore, on average, only ∼5.5% of targets were affected by *MYC* ecDNA mobile interactions. This paradigm needs to be addressed in future studies by single-cell ATAC-seq and RNA-seq to reveal the actual effect size in individual cells, and to understand whether such cell-to-cell variation may provide clonal heterogeneity and advantage for cancer development. Nonetheless, this observation suggests that, unlike HSR in the chromosome that mediates stable inheritance to ensure maximized oncogene output, ecDNA is built for variation, which is by heterogeneously *trans*-activating transcription among individual cancer cells, as well as copy number variation through random segregation^23^.

ecDNAs can aggregate and form transcription-promoting hubs ^1^. We found that ecDNA hub size varies, ranging from micron-size ecDNA speckle to small ecDNA minim made of a few ecDNA particles. Interestingly, the likelihood of ecDNA aggregation is positively correlated with ecDNA’s ability to act *in trans* with chromosomes. Given that both interactions are virtually chromatin-chromatin interactions mediated by the same ecDNA-borne nascent lncRNA, we conclude that ecDNA-chromosome *trans*-interaction and ecDNA hub formation are the two sides of the same coin.

LncRNA is an essential component of many membrane-less organelles as nuclear bodies ^24^. For example, a lncRNA, nuclear paraspeckle assembly transcript 1 (*NEAT1*), acts as a scaffold to recruit RNA binding proteins, including NONO, SFPQ, and FUS, to form paraspeckles ^25^. These paraspeckles assemble upon the transcription of *NEAT1* and usually organize closely to the *NEAT1* gene locus ^26^. FUS, as an essential paraspeckle protein, interacts with *NEAT1* through the prion-like domain, which is crucial for liquid-liquid phase separation. *NEAT1* traps specific RNA binding proteins in paraspeckle to restrict their activity in the nucleoplasm ^27^ and interacts with transcription start sites of active genes in breast cancer cells for potential transcriptional regulation^28^. We found that lncRNA *PVT1* is responsible for ecDNA speckle organization, representing a new example of DNA-centered, lncRNA-organized nuclear structure. Whether the ecDNA hub is also a type of phase separation, and therefore, creating a unique therapeutic window, remains to be further explored.

Interestingly, we found that the knockdown of *PVT1* mainly dissociated ecDNA speckle, but not ecDNA minim, although it is difficult to rule out the contribution of the residual *PVT1* in ecDNA minim. Given the evidence that the histone-binding protein BRD4 concatenates ecDNA contacts ^1^, it is possible that RNA and protein functionally couple together to drive the speckle and minim formation of ecDNA. We show that disrupting ecDNA speckles can decrease the expression of ecDNA-borne genes, suggesting that ecDNA aggregation enables enhancer hitchhiking among ecDNA molecules.

Transcriptional regulation of the oncogene *MYC* becomes more complicated when amplified, especially co-amplified with *PVT1*. The *PVT1* promoter was previously found to be a tumor suppressor element, which competes with the *MYC* promoter for functional enhancers ^22^. However, *MYC* ecDNA in COLO320DM cells utilized the *PVT1* promoter to drive *MYC* expression as a fusion gene, termed “promoter hijacking”, to achieve high transcriptional activity^22^. Moreover, we observed that the expression of promoter-hijacked *MYC* is resistant to ecDNA speckle dispersion, suggesting that gene fusion may confer plasticity of oncogene output in cancer. In addition, the circular configuration of ecDNA enables ultra-long-range chromatin interaction, which can provide additional enhancer input for an oncogene ^12^. In summary, oncogene *MYC* can be regulated by its native enhancers, the distal enhancers via ecDNA circularization, other enhancers by ecDNA aggregation, and gene fusion as an additional fail-safe mechanism. Therefore, it will be necessary to dissect the transcriptional program of an oncogene in multiple aspects, especially when it is amplified on ecDNA.

## METHODS

### Cell culture

Human gastric carcinoma cell line SNU-16, and human colon cancer lines COLO320DM and COLO320HSR were cultured in DMEM/F12 with 10% fetal bovine serum (FBS). Human glioblastoma GBM39EC and GBM39HSR tumor spheroids were cultured in DMEM/F12 with GlutaMAX, B27, EGF (20 ng/ml), FGF (20 ng/ml) and 5 μg/ml heparin.

### Metaphase chromosome spread

Cells in metaphase were generated by KaryoMAX (Gibco) treatment at 0.1 μg/ml for 3 h (COLO320) or overnight (GBM39). Single-cell suspension was obtained, washed with PBS, and treated with 75 mM KCl for 15-30 min. Samples were then fixed by Carnoy’s fixative (3:1 methanol:glacial acetic acid, v/v) and washed three times with fixative. The cell pellet in fixative was dropped onto humidified glass coverslips.

### DNA FISH

Slides containing fixed cells in metaphase were briefly equilibrated by 2× SSC buffer, followed by dehydration in 70%, 85%, and 100% ethanol for 2 min each. FISH probes (Empire Genomics) were added to the slide, and a coverslip was applied. Samples were co-denatured with probes at 75 °C for 1 min on a hotplate and hybridized at 37 °C overnight. After removing the coverslip, samples were washed once by 0.4× SSC, and twice with 2× SSC with 0.05% Tween-20 for 2 min each. DNA was stained with DAPI (1 μg/ml) for 2 min, washed with 2× SSC, and then the mounting medium was applied before imaging.

### Microscopy

DNA FISH images were acquired with Zeiss Axio Observer 7 microscope with the Apotome 3 optical sectioning module, using 63× Plan-APOchromat NA 1.4 oil lens. With the Apotome optical sectioning turned on, at least five z-stacks were taken with an interval of 1-µm. To calculate the ecDNA cluster score, only one z-stack with the highest signal of interest was chosen from each nucleus for analysis.

### ecDNA cluster score

When an ecDNA hub occurs, the pixels that indicate ecDNA particles aggregate. The pixel density peaks in the center of the cluster and decreases with distance from the center. This feature was fitted into a Gaussian distribution and described by a probability density function (PDF), which is a statistical expression to determine the likelihood of an outcome, to model ecDNA aggregation. A higher PDF indicates that pixels are more likely to aggregate. When the PDF reached a certain threshold, we defined it as the presence of an ecDNA hub. Multiple ecDNA hubs were fitted by multiple PDFs, forming a Gaussian mixture model to describe the likelihood of ecDNA aggregation.

### GRID-seq library construction

A bivalent linker was chemically synthesized, including biotin-labeled DNA strand and DNA/RNA hybrid strand: 5Phos-GTTGGAGTTCGGTGTGTGGGAG/iBiodT/GACCAGTGTC and 5Phos-rGrUrUrGrGrArCrANNNNGACACTGGTCACTCCCACACACCGAACTCCAAC. The DNA/RNA hybrid strand was pre-adenylated with DNA 5’ Adenylation Kit (NEB) and purified with Oligo Clean & Concentrator (Zymo). Two strands were mixed to 8 pmol/μl, heated at 80 °C for 5 min, and annealed by slow cooling to room temperature.

Approximately 1 million mammalian cells were collected, washed twice with 1× PBS, and crosslinked with 2 mM DSG for 45 min and 3% formaldehyde followed at room temperature for 10 min. 1.2 M Glycine was applied to quench formaldehyde at the final concentration of 350 nM, and cells were washed twice with PBS. Fixed cells were suspended in Equilibrium Buffer (10 mM Tris-Cl pH 7.5, 10 mM NaCl, 0.2% IPGAL, 0.8 U/μl RiboLock, 1× protease inhibitor) for 15 min on ice, washed with 1× Tango Buffer with 0.1% Tween-20, and incubated in SDS buffer (1× Tango Buffer, 0.4% SDS) at 62 °C for 10 min. SDS was immediately quenched with Triton X-100 at the final concentration of 1.5%, and well-isolated nuclei were collected. Nuclei were treated by AluI solution (1× Tango Buffer, 0.8 U/μl RiboLock, 1× protease inhibitor, 1% Triton X-100, 0.5 U/μl Alu I Thermo) at 37 °C for 2 h, and PNK solution (1× Tango Buffer, 0.8 U/μl RiboLock, 1 mM ATP, 10 U/μl PNK) at 37 °C for 1.5 h followed.

Nuclei were pre-washed and applied for *in situ* RNA linker ligation (1× RNA ligase buffer, 0.8 U/μl RiboLock, 100 pmol pre-adenylated linker, 4 U/μl T4 RNA ligase 2 truncated KQ, 15% PEG-8000) at 25 °C for 2 h. Linker contained primer was extended with SuperScript^TM^ III First-Strand Synthesis System Kit, and applied for dsDNA ligation (1× DNA ligase buffer w/o PEG, 0.8 U/μl RiboLock, 1 mg/ml BSA, 1% Triton X-100, 1 U/μl T4 DNA Ligase) at 16 °C overnight. Cells were treated by Proteinase K buffer (50 mM Tris-Cl pH 7.5, 100 mM NaCl, 1 mM EDTA, 1% SDS, 1 mg/ml Proteinase K) at 65 °C for 30 min, adjusted to 450 mM NaCl and then DNA/RNA was precipitated with isopropanol. RNA was replaced into DNA in RNase H solution (1× NEBuffer 2, 0.5 mM dNTP, 5 U Rnase H, 0.5 U/μl E. coli DNA pol. I) at 16 °C for 3 h, selected by 50 μg of biotin-labeled streptavidin beads, and applied for nick sealing solution (1× E. coli DNA ligase buffer, 0.15 U/μl E. coli DNA ligase) at 16 °C for 1 h. ssDNA was generated by 100 μl of 150 mM NaOH at room temperature for 10 min, neutralized with 6.5 μl of 1.25 M acetic acid, and diluted with 11 μl of 10× TE Buffer (100 mM Tris-Cl pH 7.5, 10 mM EDTA). Then non-biotin labeled ssDNA was mixed with 250 ng Random Hexamer at 55 °C for 5 min, chilled on ice, and followed second strand generation solution (1× NEBuffer 2, 15 U Klenow exo^-^) at 25 °C for 10 min and 37 °C for 40 min. dsDNA was digested by MmeI solution (1× NEB Cutsmart, 2 U MmeI) at 37 °C for 0.5 h and extracted with 12% native polyacrylamide TBE gel (∼84 bp band). Purified dsDNA was removed phosphate by 0.5 U rSAP, ligated to 125 pmol annealed NN-adaptor and extracted with 10% native polyacrylamide TBE gel (∼180 bp band). Then purified dsDNA was amplified by PCR with 13 cycles, extracted with 10% native polyacrylamide TBE gel (216 bp band) and purified for sequencing.

### GRID-seq data processing

Paired-end reads of GRID-seq were joined into continued sequence by flash software (-z -M 90 -m 20), aligned to GRID-seq linker with bwa software (-k 5 -L 4 -B 2 -O 4), and then divided into RNA (cDNA) or DNA (genomic DNA) reads according to the linker position. RNA and DNA reads were separately aligned to human genome (hg38) with bwa (-k 17 -w 1 -T 1), and PCR duplicates were removed using UMI, 4 bp sequence in GRID-seq linker. Then RNA or DNA reads were classified into unique or multiple alignments according to the MAPQ (larger than 0 or not). Unique aligned RNA or DNA reads were collected, and multiple aligned RNA or DNA reads were re-mapped to the genome with bwa (-k 17 -w 1 -T 1 -a -c 50/100 (for RNA or DNA, respectively)) to assign a weighted value of an RNA or DNA read according to the surrounded density (within 1 kb) of unique DNA or RNA read that paired. Then, all weighted RNA-DNA pairs were collected for downstream analysis.

According to the previously described strategy ^21^, the chromatin-associated RNAs (caRNAs) were identified, following specific RNA-chromatin interactions were detected against the non-specific background built by protein-coding RNAs (pcRNAs), and interaction heatmap analysis were conducted in 2 Mb resolution. RNA reads derived from amplicons (ecDNA and HSR) were isolated, and the paired DNA reads were aggregated into density of 1 kb resolution bins. Aggregated signals exceeding 2-folds of background were clustered into continuous region and further constructed an averaged signal for the region. The sub-region containing a signal higher than average one was further identified as the high confident target of ecDNA or HSR-derived RNAs.

Intronic, exonic, or overlapped regions were identified according to all the transcripts annotated by GENCODE (version 31). Using bedtools software, RNA reads overlapped to the intronic or exonic regions were identified as the source of nascent RNA or nuclear-retained RNA, respectively. Then, specific RNA-DNA interactions of intronic or exonic reads were separated to generate heatmap in 100 kb resolution.

### Hi-C library construction and data processing

Approximately 1 million mammalian cells were collected and then applied for BL-Hi-C approach as previously described ^18^ to construct the Hi-C libraries under restrict enzyme AluI-based version. The HiC-Pro software was applied to generate interacting matrix analysis with default parameters, and ChIA-PET2 software was used to detect highly confident chromatin interactions ^29^. The metric of normalized *trans*-chromosomal interaction frequency (nTIF) was carried out in 50 kb resolution of DNA bins, according to the previously described method ^2^. WGS, ATAC-seq, and RNA-seq signals were normalized (CPM), and calculated the density on the overlapped 50 kb bins with bedtools software. Adjustment of nTIF against the WGS signal of each bin in the log-2 scale was calculated and further normalized by the median value collectively from all chromosomal bins. Fold enrichment of WGS, ATAC-seq, or RNA-seq signals against nTIF in log-2 scale was calculated and further normalized by the median value collectively from all chromosomal bins, respectively.

### ATAC-seq library construction and data processing

ATAC-seq library was constructed as previously described ^30^. Adaptor was trimmed with Cutadapt software, and then reads were aligned to the hg38 genome using Bowtie2 software. Duplicates were removed with Picard software, and peaks were called using MACS2. The read counts of each sample on the peak were identified using bedtools software, and followed DESeq2 software was applied to detect differential chromatin accessibility with FDR less than 0.05.

### Antisense oligo transfection

2× 10^5^ cells were plated on a 12-well plate, and ASO was transfected into the cells using RNAiMAX immediately. PVT1 ASO was used as previously described ^22^ with the final concentration of 250 nM in 1 ml media. One day after transfection, cells were harvested for downstream libraries construction and qRT-PCR analysis.

### RT-qPCR

RNA was prepared with RNA Extraction Kit (Zymo) and quantified by Nanodrop. Then, cDNA was prepared by SuperScript^TM^ III First-Strand Synthesis System Kit was applied on 1 μg of RNA, and the CT value was measured using Applied Biosystems^TM^ StepOnePlus^TM^ Real-Time PCR System, and each mean dCT was averaged from triplicates. The relative expression level was measured by ddCT according to *GAPDH* control as previously described ^22^.

### Quantifications and Statistical Analysis

Statistical parameters were reported either in individual figures or corresponding figure legends. Quantification data are, in general, presented as bar/line plots, with the error bar representing mean ±SEM, or boxplot, showing the median (middle line), first and third quartiles (box boundaries), and furthest observation or 1.5 times of the interquartile (end of whisker). All statistical analyses were done in R. Whenever asterisks were used to indicate statistical significance, * stands for p < 0.05; ** p < 0.01, and *** p < 0.001.

### Data Availability

Data generated in this study have been deposited in NCBI SRA with accession number PRJNA989676.

### Code availability

Custom code in this work is available upon reasonable request from the corresponding authors. Software for ecDNA cluster score is deposited in GitHub (https://github.com/grapeot/ecDNAClusterScore)

## ACKNOWLEDGMENTS

We thank members of the Fu and Wu laboratories for discussions. Z.L. was supported by the startup funding of SUSTech (Y011176102, Y011176202). S.W. is supported by the Cancer Prevention and Research Institute of Texas (RR210034) and Gilead’s Research Scholars Program in Solid Tumors. This work was delivered as part of the eDyNAmiC team supported by the Cancer Grand Challenges partnership funded by Cancer Research UK (P.S.M CGCATF-2021/100012, S.W. CGCATF-2021/100023) and the National Cancer Institute (P.S.M. OT2CA278688, S.W. OT2CA278683).

## AUTHOR CONTRIBUTIONS

Z.L., S.W., and X.-D.F. conceived the project. Z.L. performed the majority of experiments and most genomic data analysis. C.G., W.L., Y.W., and S.W. contributed to imaging experiments and data analysis. Y.W. developed the method for ecDNA clustering score. Z.L. and S.W. prepared figures. M.Q.Z. and D.-E.Z. contributed to data interpretation. Z.L., S.W., and X.-D.F. wrote the manuscript.

## SUPPLEMENTAL VIDEO, DATA, AND EXCEL TABLE TITLES AND LEGENDS

**Supplementary Table 1:** All of oligos used are listed, including ASO, primers and linkers.

**Supplementary Table 2:** All regents used for the libraries construction are listed.

**Supplementary Table 3:** All softwares used for data analysis are listed.

**Supplementary Table 4:** GRID-seq data and optimization of multiple alignment.

---

**Extended Data Figure 1.**
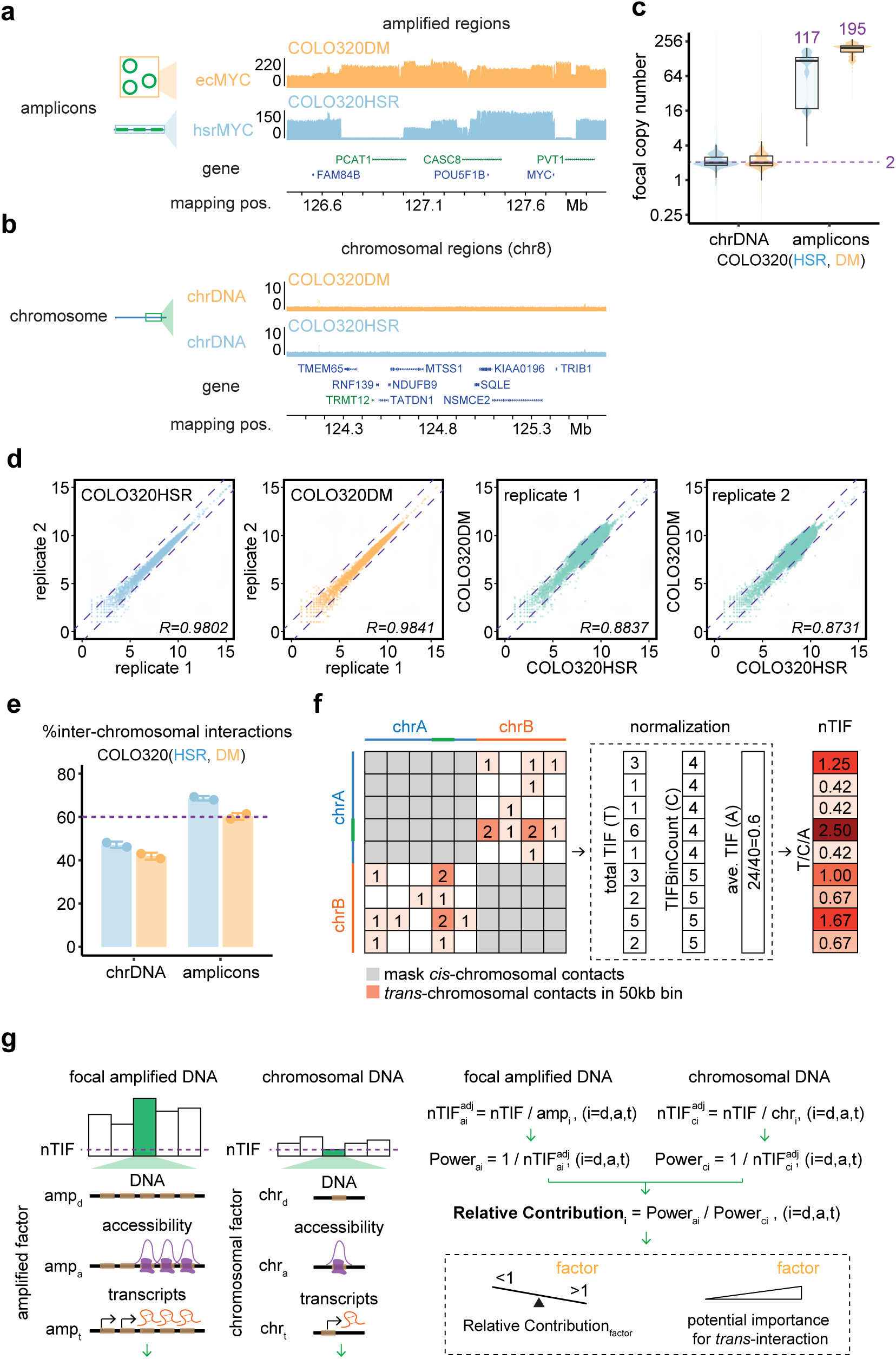
Focal amplicons of ecDNA and *in situ* Hi-C detected contacts in COLO320 cells, Related to Figure 1. **(a-b)** WGS tracks for focal amplified ecMYC (orange) and hsrMYC (blue) in COLO320 cells (a) and WGS tracks for representative chromosome 8 region as an internal control (b). **(c)** WGS estimated copy numbers for chromosomal regions and focal amplificons of hsrMYC or ecMYC in COLO320 cells. **(d)** Spearman correlation of Hi-C libraries between replicates (left two) or conditions (right two) in 50-kb resolution. **(e)** Percent of inter-chromosomal read pairs detected in Hi-C assay for chromosomal regions or amplified regions in COLO320 cells. **(f)** Schematic of normalized *trans-*chromosomal interaction frequency (nTIF). *Trans-* chromosomal interactions between DNA bins (50-kb resolution) were used and normalized to the number of interacting bins and averaged nTIF. **(g)** Schematic of relative enrichment analysis to estimate amplicon-associated factors linked to the interactions *in trans*. Genomic segments carried factors of copy number (WGS, amp_d_), chromatin accessibility (ATAC-seq signal, amp_a_), or transcripts (RNA-seq signal, amp_t_) were used to normalize those segments-derived nTIF (nTIF^adj^). The adjusted nTIF was converted to the normalization power of each factors for amplicons (Power_ai_) and chromosomal regions (Power_ci_). Relative contribution was calculated by the comparison between amplicons and chromosomal regions.

**Extended Data Figure 2.**
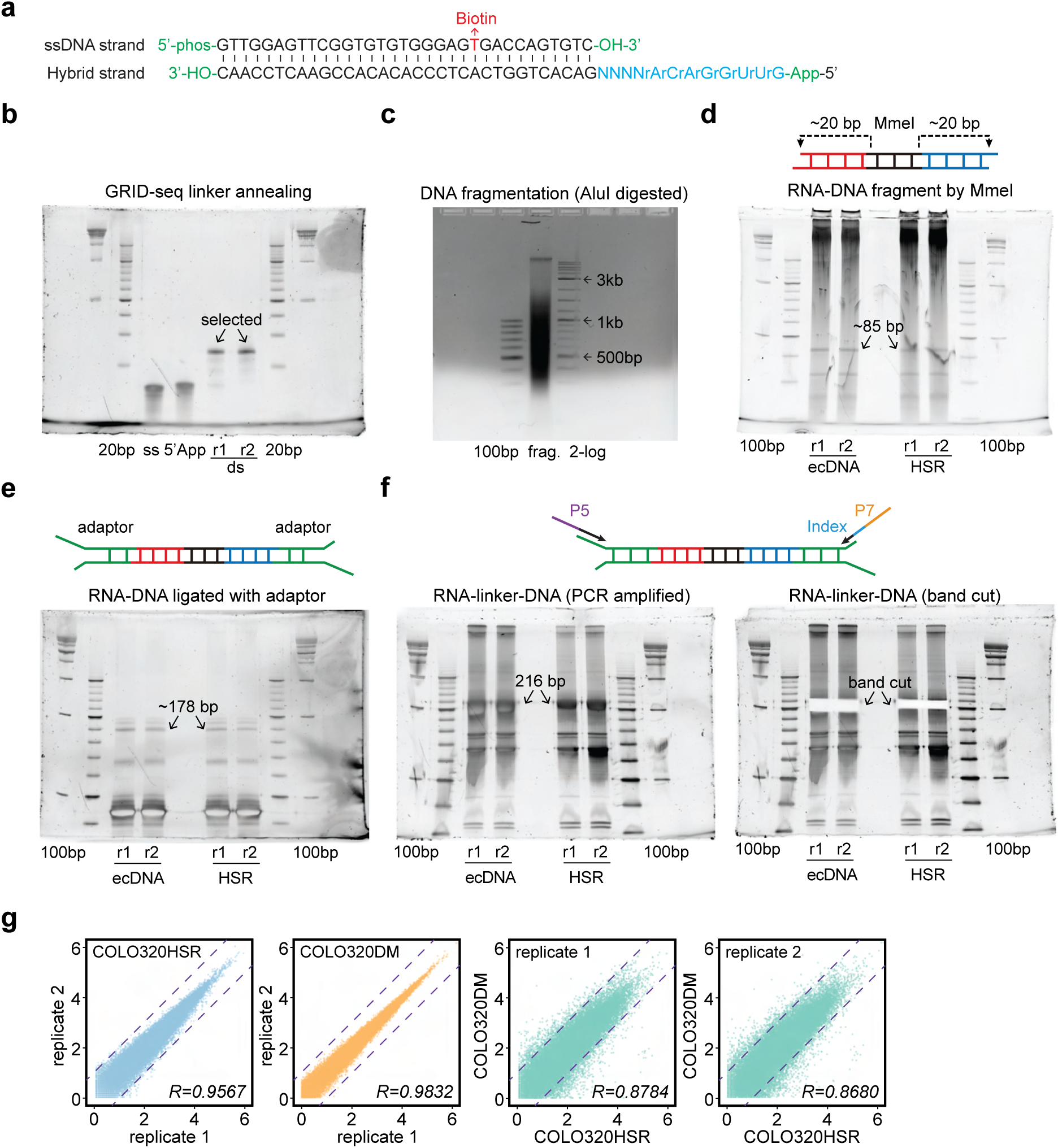
GRID-seq libraries of COLO320 cells, Related to Figure 2. **(a)** The bivalent linker in GRID-seq. App, pre-adenylation. **(b-c)** Polyacrylamide gel image of GRID-seq linker annealing (b) and chromatin fragmentation by restriction enzyme Alu I (c). **(d-f)** Polyacrylamide gel image of library products after paired-end tagging by MmeI (85 bp, d), adaptor ligation (∼178 bp, e), and PCR amplification (∼216 bp, f). Arrows indicate the DNA bands to be recovered. **(g)** Spearman correlation of GRID-seq libraries between replicates (left two) or conditions (right two) in 50-kb resolution.

**Extended Data Figure 3.**
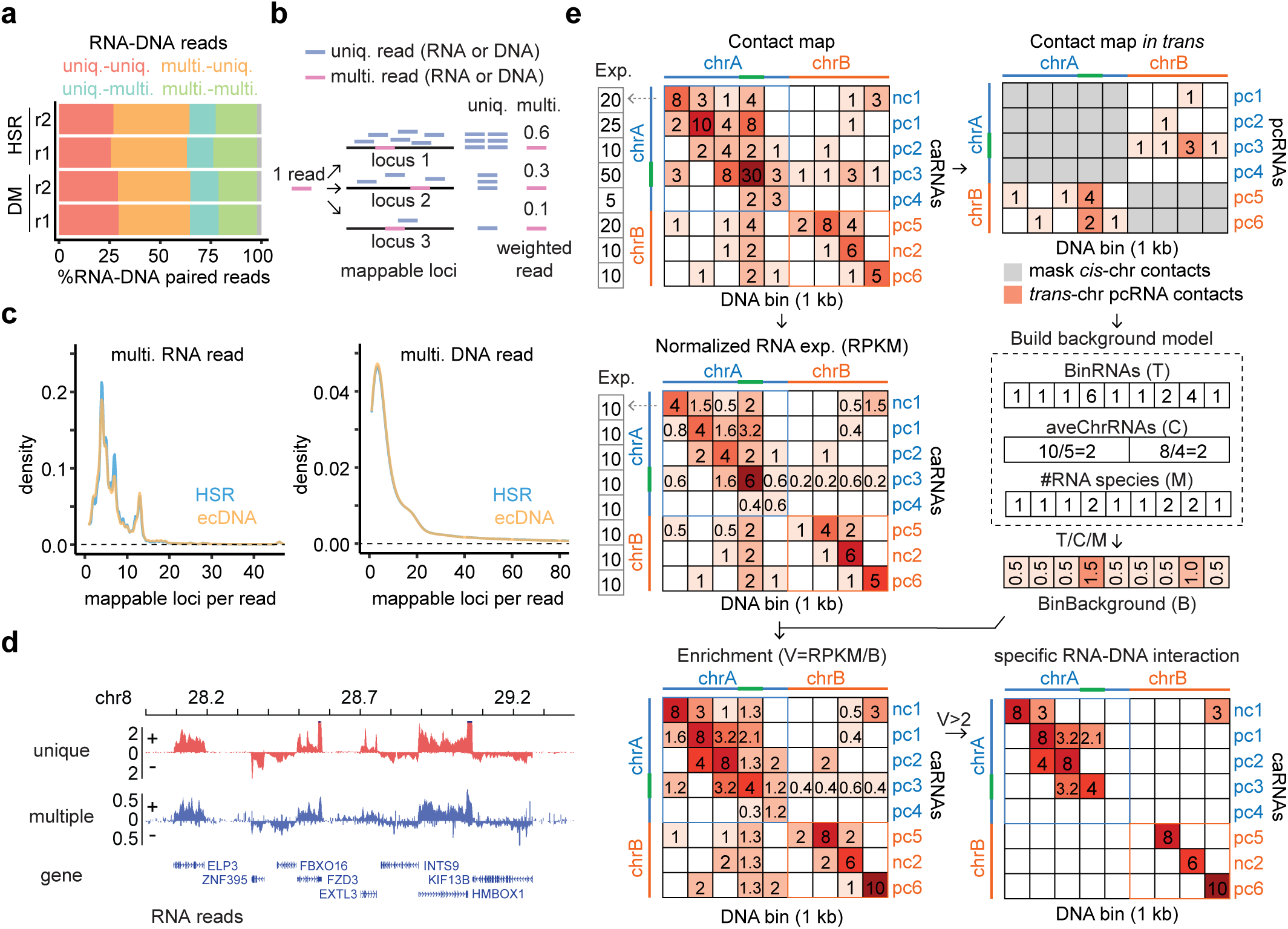
GRID-seq data analysis, Related to Figure 2. **(a)** Percent of GRID-seq assay detected each RNA or DNA read mapping to the human genome uniquely for one locus (uniq) or multiple for several loci (multi). **(b)** Weighted strategy for each multiple aligned read by estimating surrounding uniquely aligned reads where the reads were potentially mapped. **(c)** Density distribution of the locus number where multiple aligned RNA reads (left) or DNA reads (right) were mapped. **(d)** Representative genomic loci showing RNA signals derived from uniquely aligned RNA reads (red) or multiple aligned RNA reads with weighted strategy (blue). **(e)** Procedure of GRID-seq data processing. Chromatin-enriched RNAs interacting with DNA bins (1-kb resolution) were built into a matrix (top-left). Each chromatin-enriched RNA-associated interaction was normalized to the RNA expression as RPKM (middle-left). Protein-coding RNAs (pc) were gathered to build the background model by normalization to the bin number, chromosome length, and RNA species (top-right and middle-right). Relative fold enrichment was calculated for the RNA-DNA contacts against the background model of the interacted DNA bin (bottom-left), and specific RNA-DNA interaction was identified with relative fold enrichment large than 2 (bottom-right).

**Extended Data Figure 4.**
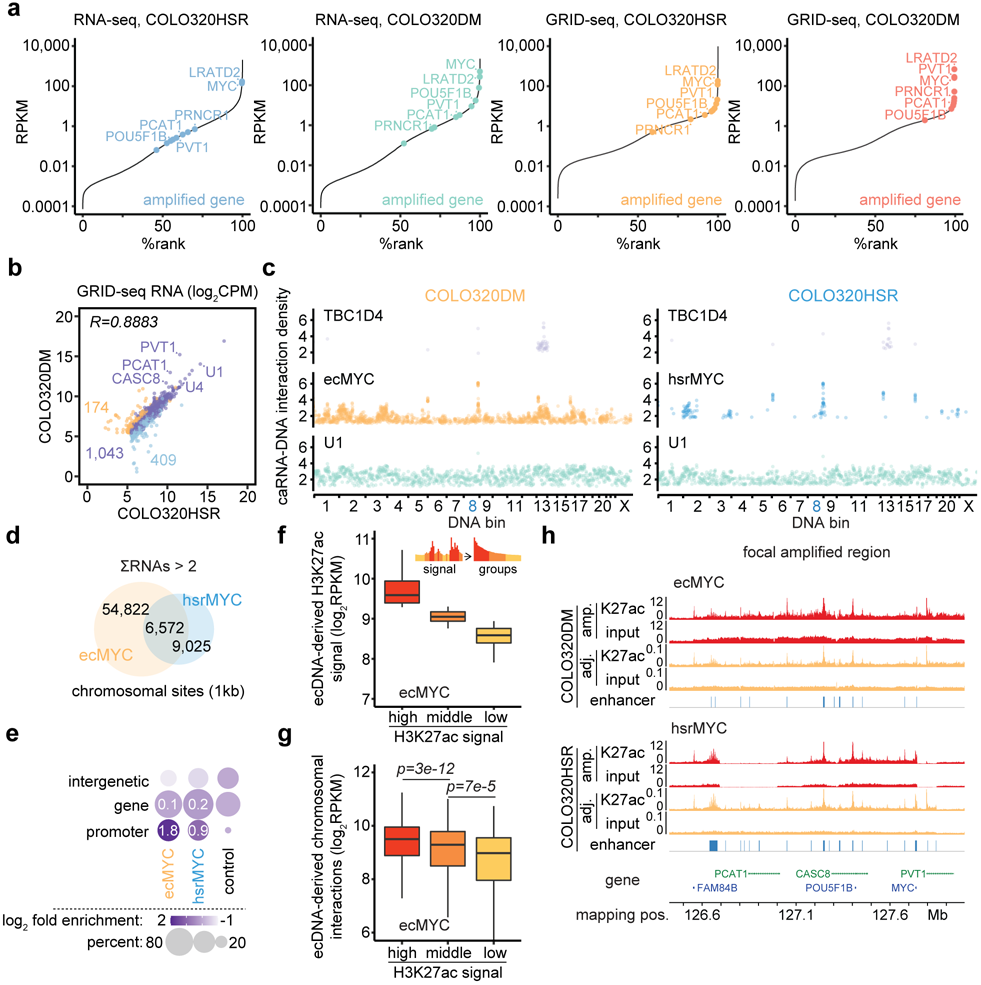
ecDNA enhancers target chromosomal promoters in COLO320DM cells, Related to Figure 2. **(a)** Gene expression within the transcriptome of COLO320HSR or COLO320DM cells detected by RNA-seq assay (left two) or GRID-seq assay (right two). Dots denote ecDNA or HSR-derived genes. **(b)** Scatter plot of RNA expression detected by GRID-seq in COLO320HSR cells against COLO320DM cells. Numbers indicate chromatin-enriched RNAs detected specifically in COLO320HSR cells (blue), COLO320DM cells (orange), or shared (purple). Spearman correlation was applied. **(c)** Genomic positions interacted by representative chromatin-enriched RNA, e.g., protein-coding RNA *TBC1D4*, ecMYC or hsrMYC amplified RNA *PVT1*, and well-known *trans-*functioning spliceosome RNA *U1* in COLO320DM and COLO320HSR cells. **(d)** A Venn diagram showing the number of chromosomal loci (1-kb resolution) interacted by ecMYC or hsrMYC-derived RNAs in COLO320DM or COLO320HSR cells. **(e)** Genomic annotation for chromosomal targets of ecMYC (orange), hsrMYC (blue), or randomly shuffled chromosomal segments (nc, black), with log-2 scaled relative enrichment in ranged color and percentage in dot size. **(f-g)** Quantification of H3K27ac signal (f) and Hi-C interactions *in trans* (g) stratified by the H3K27ac signal in ecMYC regions with 1-kb bins (one-sided Wilcoxon Rank Sum Test). **(h)** Representative genomic loci showing H3K27ac ChIP-seq signal (red) and copy number adjusted H3K27ac ChIP-seq signal (orange) for amplified regions in COLO320DM and COLO320HSR cells. Enhancers were called by the ROSE software using copy number adjusted H3K27ac signal.

**Extended Data Figure 5.**
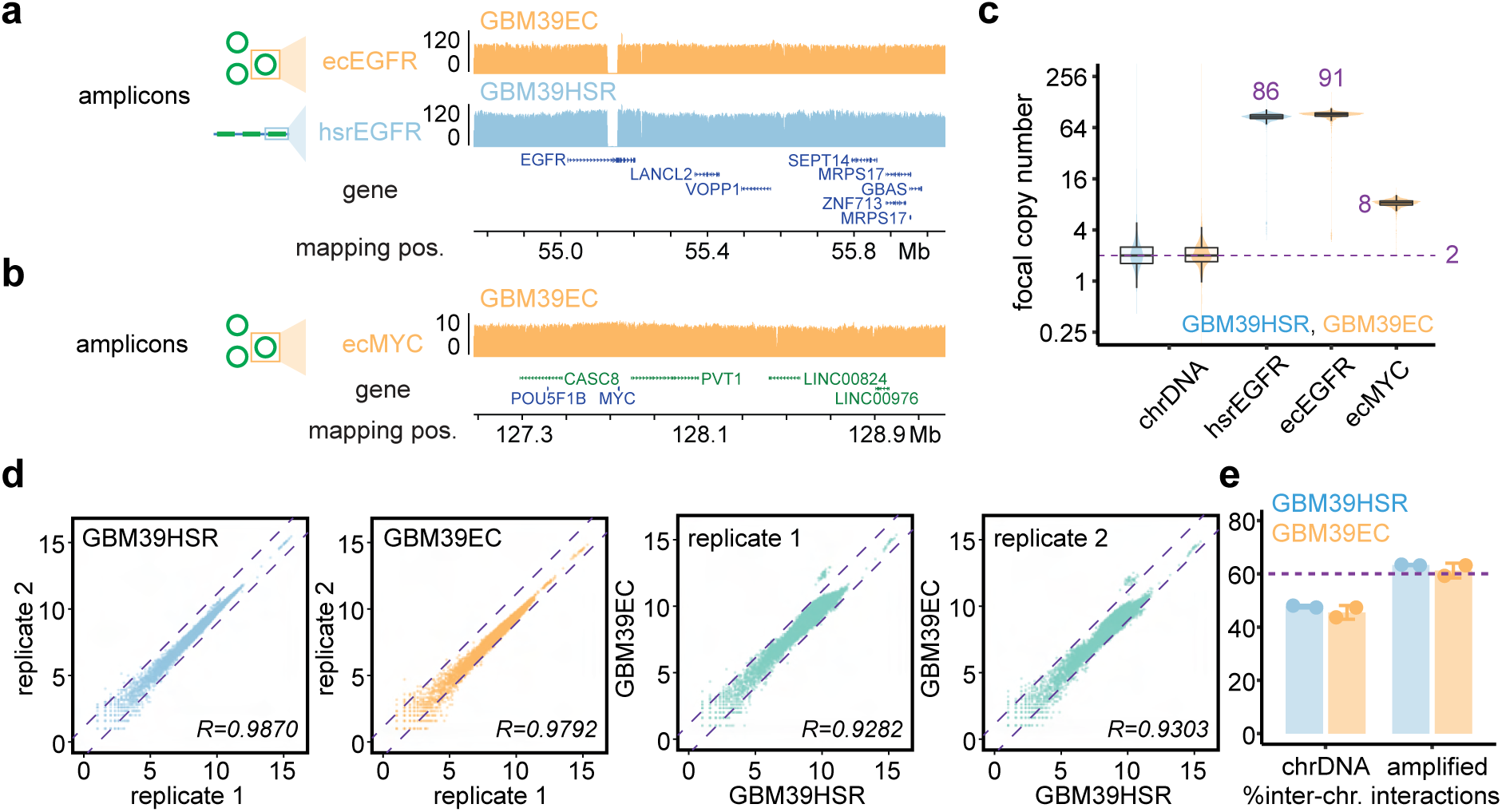
Focal amplicons of ecDNA and in situ Hi-C detected contacts in GBM39 cells, Related to Figure 3. **(a-b)** WGS tracks for focally amplified ecEGFR (orange) and hsrEGFR (blue) in GBM39 cells (a) and ecMYC (orange) in GBM39EC cells (b). **(c)** WGS estimated copy numbers for chromosomal regions and amplicons of hsrEGFR, ecEGFR, and ecMYC in GBM39 cells. **(d)** Spearman correlation of Hi-C libraries between replicates (left two) or conditions (right two) in 50-kb resolution. **(e)** Percent of inter-chromosomal read pairs detected in Hi-C assay for chromosomal regions or amplified regions in GBM39 cells.

**Extended Data Figure 6.**
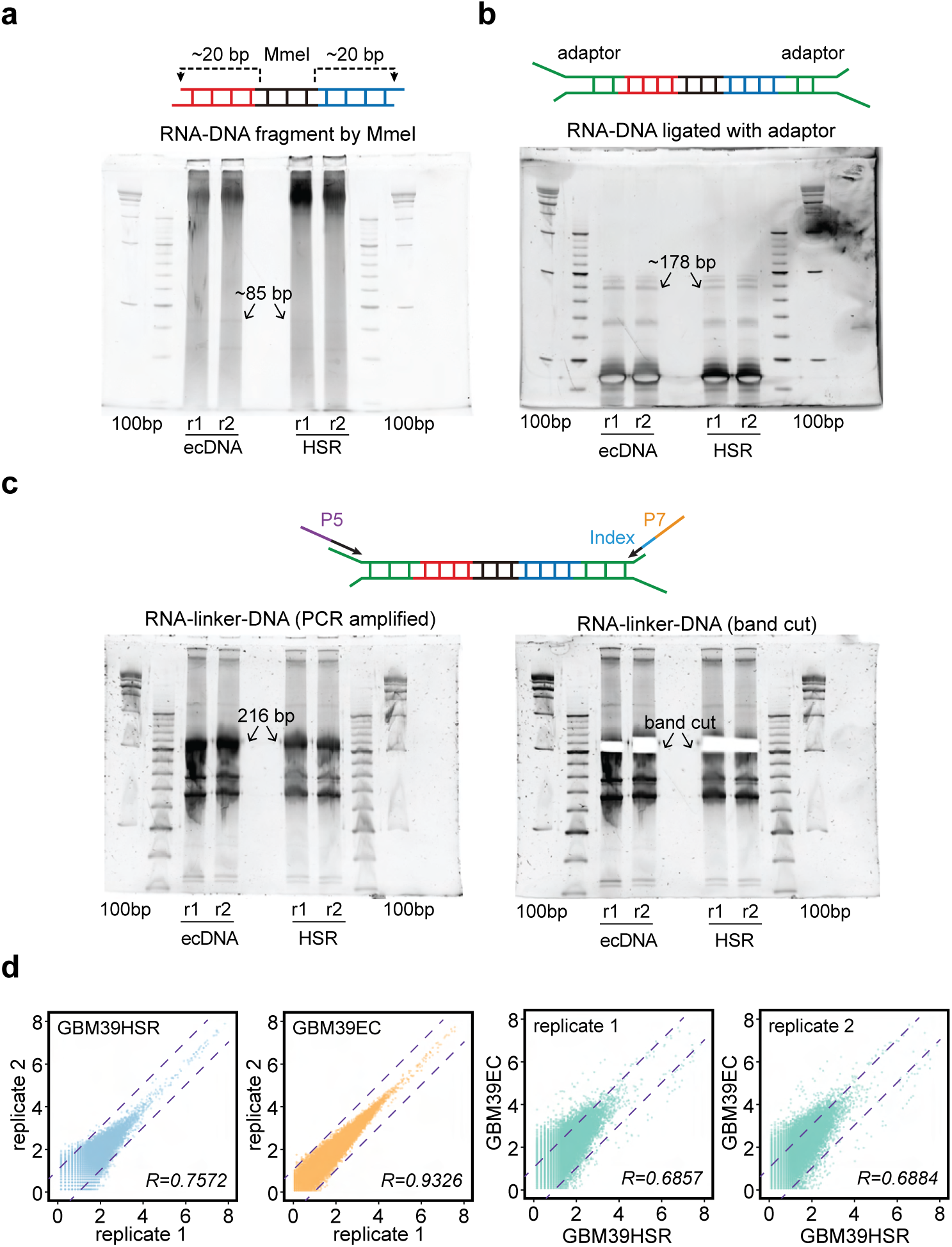
GRID-seq libraries of GBM39 cells, Related to Figure 3. **(a-c)** Polyacrylamide gel image of library products after paired-end tagging by MmeI (85 bp, a), adaptor ligation (∼178 bp, b), and PCR amplification (∼216 bp, c). Arrows indicate the DNA bands to be recovered. **(d)** Spearman correlation of GRID-seq libraries between replicates (left two) or conditions (right two) in 50-kb resolution.

**Extended Data Figure 7.**
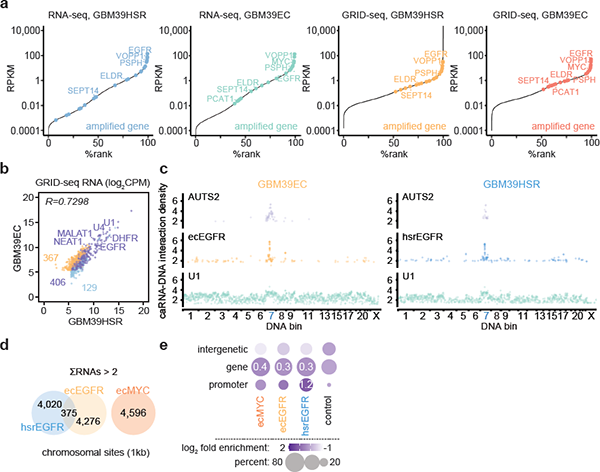
ecDNA mobile targeting analysis in GBM39 cells, Related to Figure 3. **(a)** Gene expression within the transcriptome of GBM39HSR and GBM39EC cells detected by RNA-seq assay (left two) or GRID-seq assay (right two). Dots denote amplicon-derived genes. **(b)** Scatter plot of RNA expression detected by GRID-seq in GBM39HSR cells against GBM39EC cells. Numbers indicate chromatin-enriched RNAs detected specifically in GBM39HSR cells (blue), GBM39EC cells (orange), or shared (purple). Spearman correlation was applied. **(c)** Genomic positions interacted by representative chromatin-enriched RNA, e.g., protein-coding RNA *AUTS2*, ecEGFR or hsrEGFR amplified RNA *EGFR*, and well-known *trans-*functioning spliceosome RNA *U1* in GBM39EC and GBM39HSR cells. **(d)** A Venn diagram showing the number of chromosomal loci (1-kb resolution) interacted by ecEGFR, ecMYC, or hsrEGFR-derived RNAs in GBM39EC or GBM39HSR cells. **(e)** Genomic annotation for chromosomal targets of ecMYC, ecEGFR, hsrEGFR, or randomly shuffled chromosomal segments (nc, black), with log-2 scaled relative enrichment in ranged color and percentage in dot size.

**Extended Data Figure 8.**
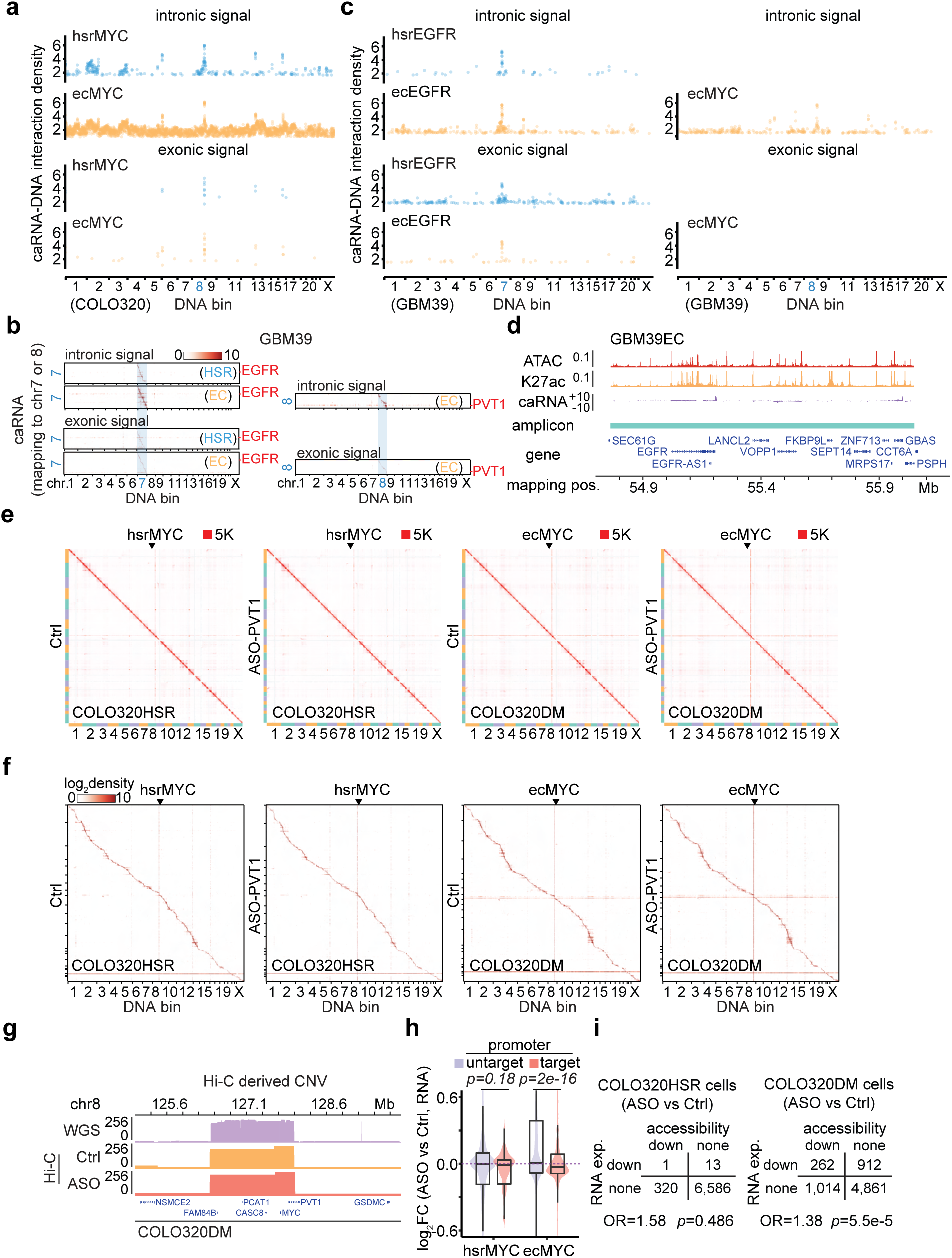
Characterization of ecDNA-borne ncRNAs for ecDNA interactome, Related to Figure 4. **(a)** Genomic positions interacted by hsrMYC or ecMYC-derived intronic signal (left) or exonic signal *MYC* (right) in COLO320 cells. **(b)** Heatmaps of hsrEGFR, ecEGFR, or ecMYC-derived RNAs interacting with the genome in the order of mapping position in GBM39 cells. Intronic and exonic RNA signals were used to generate the heatmaps independently. Scale bars, log-10 relative enrichment. **(c)** Genomic positions interacted by hsrEGFR or ecEGFR-derived intronic signal or exonic signal in GBM39 cells (left), and ecMYC-derived intronic signal and exonic signal on right. **(d)** Genomic tracks showing ATAC-seq, H3K27ac ChIP-seq, and GRID-seq-derived RNA (strand-specific, in two scaled views) signals normalized to CPM in GBM39EC cells. Amplicons were determined by AmpliconArchitect software. **(e)** Hi-C heatmap of COLO320HSR cells (left two) and COLO320DM cells (right two) treated by control or *PVT1* knockdown, with ecMYC or hsrMYC labeled. Scale bars, 5,000 interacting reads. **(f)** GRID-seq heatmap of COLO320HSR cells (left two) and COLO320DM cells (right two) treated by control or *PVT1* knockdown with ecMYC or hsrMYC labeled. Scale bars, log-10 relative enrichment. **(g)** Representative genomic loci showing copy number of ecMYC amplicon deconvoluted by WGS or Hi-C interacting frequency. **(h)** Quantification of log-2 fold change of chromosomal genes expression upon *PVT1* knockdown against control measured by GRID-seq-derived RNA signal. Red, genes with promoters targeted by HSR or ecDNA; Purple, untargeted genes (one-sided Wilcoxon Rank Sum Test). **(i)** Odds ratio of *PVT1*-knockdown reduced chromatin accessibility determined by ATAC-seq on transcriptional responsive (down) compared to transcriptional unresponsive (none) in both COLO320HSR (left) and COLO320DM cells (right). P-values were calculated by Fisher’s exact test.

**Extended Data Figure 9.**
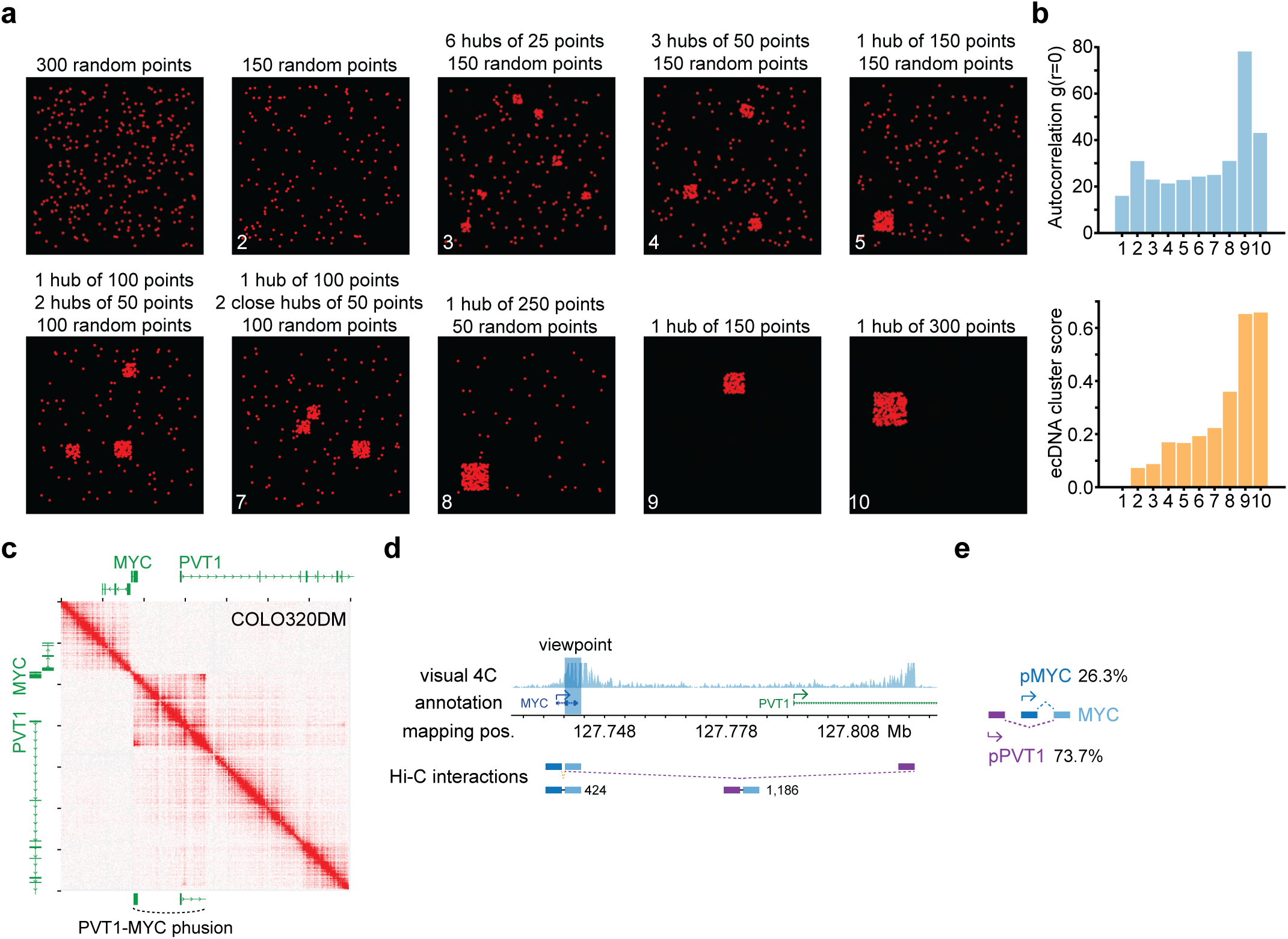
ecDNA clustering score and fusion events of *MYC* ecDNA, Related to Figure 5. **(a-b)** In silico generated pictures with points spreads, ranking by the clustering degree manually (a). Both autocorrelation g(r=0) (top) and ecDNA cluster score (bottom) were applied for the in silico generated pictures (b). **(c)** *In situ* Hi-C contact heatmap for COLO320DM cells with annotation of gene *MYC* and *PVT1* included. Fusion gene *PVT1-MYC* was illustrated. **(d-e)** Visual-4C with viewpoint of oncogene *MYC* (including exon2-3 and intron 2 in blue rectangle). Equidistant MYC promoter (4-kb in orange) and fusion PVT1 intron (4-kb in purple) derived contacts were measured (d) and were used to estimate transcript percentage (e).

## REFERENCES

1 Hung, K. L. et al. ecDNA hubs drive cooperative intermolecular oncogene expression. Nature 600, 731–736, doi:10.1038/s41586-021-04116-8 (2021).

2 Zhu, Y. et al. Oncogenic extrachromosomal DNA functions as mobile enhancers to globally amplify chromosomal transcription. Cancer Cell 39, 694–707 e697, doi:10.1016/j.ccell.2021.03.006 (2021).

3 Cremer, T. & Cremer, C. Chromosome territories, nuclear architecture and gene regulation in mammalian cells. Nat Rev Genet 2, 292–301, doi:10.1038/35066075 (2001).

4 Saurin, A. J. et al. The human polycomb group complex associates with pericentromeric heterochromatin to form a novel nuclear domain. J Cell Biol 142, 887–898, doi:10.1083/jcb.142.4.887 (1998).

5 Franke, M. et al. Formation of new chromatin domains determines pathogenicity of genomic duplications. Nature 538, 265–269, doi:10.1038/nature19800 (2016).

6 Dixon, J. R. et al. Topological domains in mammalian genomes identified by analysis of chromatin interactions. Nature 485, 376–380, doi:10.1038/nature11082 (2012).

7 Lupianez, D. G. et al. Disruptions of topological chromatin domains cause pathogenic rewiring of gene-enhancer interactions. Cell 161, 1012–1025, doi:10.1016/j.cell.2015.04.004 (2015).

8 Maass, P. G., Barutcu, A. R. & Rinn, J. L. Interchromosomal interactions: A genomic love story of kissing chromosomes. J Cell Biol 218, 27–38, doi:10.1083/jcb.201806052 (2019).

9 Xu, Z. et al. Structural variants drive context-dependent oncogene activation in cancer. Nature 612, 564–572, doi:10.1038/s41586-022-05504-4 (2022).

10 Kim, H. et al. Extrachromosomal DNA is associated with oncogene amplification and poor outcome across multiple cancers. Nat Genet 52, 891–897, doi:10.1038/s41588-020-0678-2 (2020).

11 Wu, S., Bafna, V., Chang, H. Y. & Mischel, P. S. Extrachromosomal DNA: An Emerging Hallmark in Human Cancer. Annu Rev Pathol 17, 367–386, doi:10.1146/annurev-pathmechdis-051821-114223 (2022).

12 Wu, S. et al. Circular ecDNA promotes accessible chromatin and high oncogene expression. Nature 575, 699–703, doi:10.1038/s41586-019-1763-5 (2019).

13 Morton, A. R. et al. Functional Enhancers Shape Extrachromosomal Oncogene Amplifications. Cell 179, 1330–1341 e1313, doi:10.1016/j.cell.2019.10.039 (2019).

14 Turner, K. M. et al. Extrachromosomal oncogene amplification drives tumour evolution and genetic heterogeneity. Nature 543, 122–125, doi:10.1038/nature21356 (2017).

15 Weiser, N. E., Hung, K. L. & Chang, H. Y. Oncogene Convergence in Extrachromosomal DNA Hubs. Cancer Discov 12, 1195–1198, doi:10.1158/2159-8290.CD-22-0076 (2022).

16 Adelman, K. & Martin, B. J. E. ecDNA party bus: Bringing the enhancer to you. Mol Cell 81, 1866–1867, doi:10.1016/j.molcel.2021.04.017 (2021).

17 Purshouse, K. et al. Oncogene expression from extrachromosomal DNA is driven by copy number amplification and does not require spatial clustering in glioblastoma stem cells. Elife 11, doi:10.7554/eLife.80207 (2022).

18 Liang, Z. et al. BL-Hi-C is an efficient and sensitive approach for capturing structural and regulatory chromatin interactions. Nat Commun 8, 1622, doi:10.1038/s41467-017-01754-3 (2017).

19 Rao, S. S. et al. A 3D map of the human genome at kilobase resolution reveals principles of chromatin looping. Cell 159, 1665–1680, doi:10.1016/j.cell.2014.11.021 (2014).

20 Zhou, B. et al. GRID-seq for comprehensive analysis of global RNA-chromatin interactions. Nat Protoc 14, 2036–2068, doi:10.1038/s41596-019-0172-4 (2019).

21 Li, X. et al. GRID-seq reveals the global RNA-chromatin interactome. Nat Biotechnol 35, 940–950, doi:10.1038/nbt.3968 (2017).

22 Cho, S. W. et al. Promoter of lncRNA Gene PVT1 Is a Tumor-Suppressor DNA Boundary Element. Cell 173, 1398–1412 e1322, doi:10.1016/j.cell.2018.03.068 (2018).

23 Lange, J. T. et al. The evolutionary dynamics of extrachromosomal DNA in human cancers. Nat Genet 54, 1527–1533, doi:10.1038/s41588-022-01177-x (2022).

24 Fox, A. H., Nakagawa, S., Hirose, T. & Bond, C. S. Paraspeckles: Where Long Noncoding RNA Meets Phase Separation. Trends Biochem Sci 43, 124–135, doi:10.1016/j.tibs.2017.12.001 (2018).

25 West, J. A. et al. Structural, super-resolution microscopy analysis of paraspeckle nuclear body organization. J Cell Biol 214, 817–830, doi:10.1083/jcb.201601071 (2016).

26 Clemson, C. M. et al. An architectural role for a nuclear noncoding RNA: NEAT1 RNA is essential for the structure of paraspeckles. Mol Cell 33, 717–726, doi:10.1016/j.molcel.2009.01.026 (2009).

27 Imamura, K. et al. Long Noncoding RNA NEAT1-Dependent SFPQ Relocation from Promoter Region to Paraspeckle Mediates IL8 Expression upon Immune Stimuli. Mol Cell 54, 1055, doi:10.1016/j.molcel.2014.06.013 (2014).

28 West, J. A. et al. The long noncoding RNAs NEAT1 and MALAT1 bind active chromatin sites. Mol Cell 55, 791–802, doi:10.1016/j.molcel.2014.07.012 (2014).

29 Li, G., Chen, Y., Snyder, M. P. & Zhang, M. Q. ChIA-PET2: a versatile and flexible pipeline for ChIA-PET data analysis. Nucleic Acids Res 45, e4, doi:10.1093/nar/gkw809 (2017).

30 Corces, M. R. et al. An improved ATAC-seq protocol reduces background and enables interrogation of frozen tissues. Nat Methods 14, 959–962, doi:10.1038/nmeth.4396 (2017).

